# Functional Neuroligin-2-MDGA1 interactions differentially regulate synaptic GABA_A_Rs and cytosolic gephyrin aggregation

**DOI:** 10.1101/2022.08.08.503083

**Authors:** Tommaso Zeppillo, Heba Ali, Sally Wenger, Francisco J. Lopez Murcia, Erinn Gideons, Janetti Signorelli, Michael J. Schmeisser, Jens Wiltfang, JeongSeop Rhee, Nils Brose, Holger Taschenberger, Dilja Krueger-Burg

**Author notes:** Correspondence should be addressed to Dilja Krueger-Burg. Department of Pathology and Experimental Therapy, Institute of Neurosciences, University of Barcelona, and Bellvitge Biomedical Research Institute (IDIBELL), 08907 L’Hospitalet de Llobregat, Barcelona, Spain (Francisco J. Lopez Murcia). Tommaso Zeppillo and Heba Ali contributed equally to this work.

## Abstract

The function of GABAergic synapses is critically shaped by cell adhesion proteins that recruit GABA_A_Rs to synapses and mediate transsynaptic signalling, but the synapse-type-specific function of such synaptic adhesion proteins and their mutual interaction remain incompletely understood. A ubiquitous cell adhesion protein at GABAergic synapses is Neuroligin-2 (Nlgn2), which recruits synaptic GABA_A_Rs by promoting the assembly of the postsynaptic gephyrin scaffold. While Nlgn2 is present at virtually all GABAergic synapses throughout the forebrain, its loss affects different GABAergic synapse subtypes with different severity, indicating that synapse-specific interactors and synapse-organizer-redundancies define the function of Nlgn2 for a given synapse type. Here we investigated how Nlgn2 function at GABAergic synapses in mouse hippocampal area CA1 is modulated by two recently identified interaction partners, MDGA1 and MDGA2. We show that Nlgn2 and MDGA1 colocalize most prominently in the stratum radiatum (S.R.) of area CA1, and that combined Nlgn2 and MDGA1 deletion causes a layer-specific exacerbation of the loss of gephyrin puncta in layer S.R. seen following Nlgn2 deletion. Intriguingly, combined Nlgn2 and MDGA1 deletion concurrently ameliorates the abnormal cytosolic gephyrin aggregation, the reduction in inhibitory synaptic transmission and the exacerbated anxiety-related behavior seen in Nlgn2 knockout (KO) mice. In contrast, heterozygous deletion of MDGA2 in Nlgn2 KO mice has only minor effects on gephyrin and GABA_A_R puncta and does not normalize cytosolic gephyrin aggregates, inhibitory synaptic transmission or anxiety-related behavior. Our data indicate that MDGA1, but not MDGA2, modulates Nlgn2 function, primarily by regulating the formation of cytosolic gephyrin aggregates. Given that both Nlgn2 and the MDGA family of proteins have been linked to psychiatric disorders, such as autism and schizophrenia, our data lead to the notion that abnormal gephyrin aggregation may contribute to the pathophysiology of these disorders, and that intervention with gephyrin aggregation could present a novel therapeutic strategy.

## Introduction

Information flow in the brain is critically shaped by synapses, specialized contact sites between neurons that are not mere passive relays but actively contribute to information processing and network output. Accordingly, the molecular machinery that mediates synaptic connectivity and neurotransmission plays a central role in regulating cognition and behavior, and alterations in this machinery, e.g. due to genetic or environmental causes, feature prominently in the pathophysiology of psychiatric and neurodevelopmental disorders (1-5). Therefore, defining the molecular logic that governs the establishment and maintenance of synaptic function represents a problem of utmost importance in the development of new therapeutic strategies for these disorders.

Of particular interest in this respect are alterations in the function of γ-aminobutyric acidergic (GABAergic) inhibitory neurons and synapses, which contribute to a plethora of computational network processes in health and disease. Fast GABAergic neurotransmission is mediated by the binding of GABA to postsynaptic GABA_A_ receptors (GABA_A_Rs), whose synaptic localization and function is regulated by a complex machinery of scaffolding and cell adhesion proteins (5, 6). A key component of this GABAergic postsynaptic protein machinery is the cell adhesion protein Neuroligin-2 (Nlgn2), which binds transsynaptically to presynaptic neurexins (Nrxns) and intracellularly to gephyrin and collybistin to establish and regulate GABAergic synaptic transmission (7). Accordingly, deletion of Nlgn2 in mice results in a loss of gephyrin and GABA_A_R subunits from postsynaptic sites, and in a reduction in the frequency and/or amplitude of miniature inhibitory postsynaptic currents (mIPSCs) (7-13).

An intriguing and as yet unexplained feature of Nlgn2 function at GABAergic synapses is its apparent synapse subtype specificity. Despite being present at virtually all inhibitory synapses in the brain, deletion of Nlgn2 in mice selectively affects only a distinct subset of GABAergic synapses, most notably perisomatic, likely parvalbumin-positive synapses onto principal neurons (8-10, 12). Given that GABAergic neurons are highly diverse, with different subtypes playing distinct roles in shaping network function and behavioral output (14-18), understanding the molecular mechanisms underlying the striking synapse-type specificity of Nlgn2 function is key for elucidating its role in network function and information processing in health and disease. A likely explanation lies in the differential interaction of Nlgn2 with other synaptic adhesion proteins that differentially localize to GABAergic synapse subtypes. In recent years, multiple new organizer proteins were identified at GABAergic synapses, but the mechanisms by which these might contribute to the differential formation and function of GABAergic synapse subtypes are still largely unknown (5, 19).

Among the newly described interaction partners of Nlgn2 are the MAM-domain containing glycosylphosphatidylinositol anchor (MDGA) family proteins MDGA1 and MDGA2 (20, 21). MDGAs were proposed to negatively regulate Nlgn2 function by blocking the interaction between Nlgn2 and its presynaptic partner Nrxn, thereby potentially impairing the assembly of GABAergic synapses (21-28) (Figure 1A). Intriguingly, MDGA1 was recently shown to selectively regulate the formation of GABAergic synapses onto distal dendrites, but not proximal dendrites or somata, of hippocampal area CA1 pyramidal neurons through an interaction with presynaptic amyloid precursor protein (APP) (29). These findings support the notion that MDGAs can contribute to the diversity of GABAergic synapse function, but whether they also differentially modulate Nlgn2 function at different synapse subtypes remains completely unknown. Moreover, there is substantial controversy over whether MDGA1 (24, 25) or MDGA2 (30) - or neither (31) - are the most relevant MDGAs for Nlgn2 regulation at GABAergic synapses. We addressed these questions by investigating the function and layer-specific composition of GABAergic synapses in hippocampal area CA1 of Nlgn2 / MDGA1 double KO and Nlgn2 KO / MDGA2 heterozygous KO mice. Given that variants of both Nlgn2 (7, 32-37) and the MDGAs (20, 38-41) have been linked to schizophrenia, autism spectrum disorders, and other brain disorders, our findings have important implications not only for understanding the basic biology of GABAergic synapses, but also for identifying potential new targets for neuropsychiatric disorders linked to dysfunction of GABAergic inhibition.

**Figure 1.**
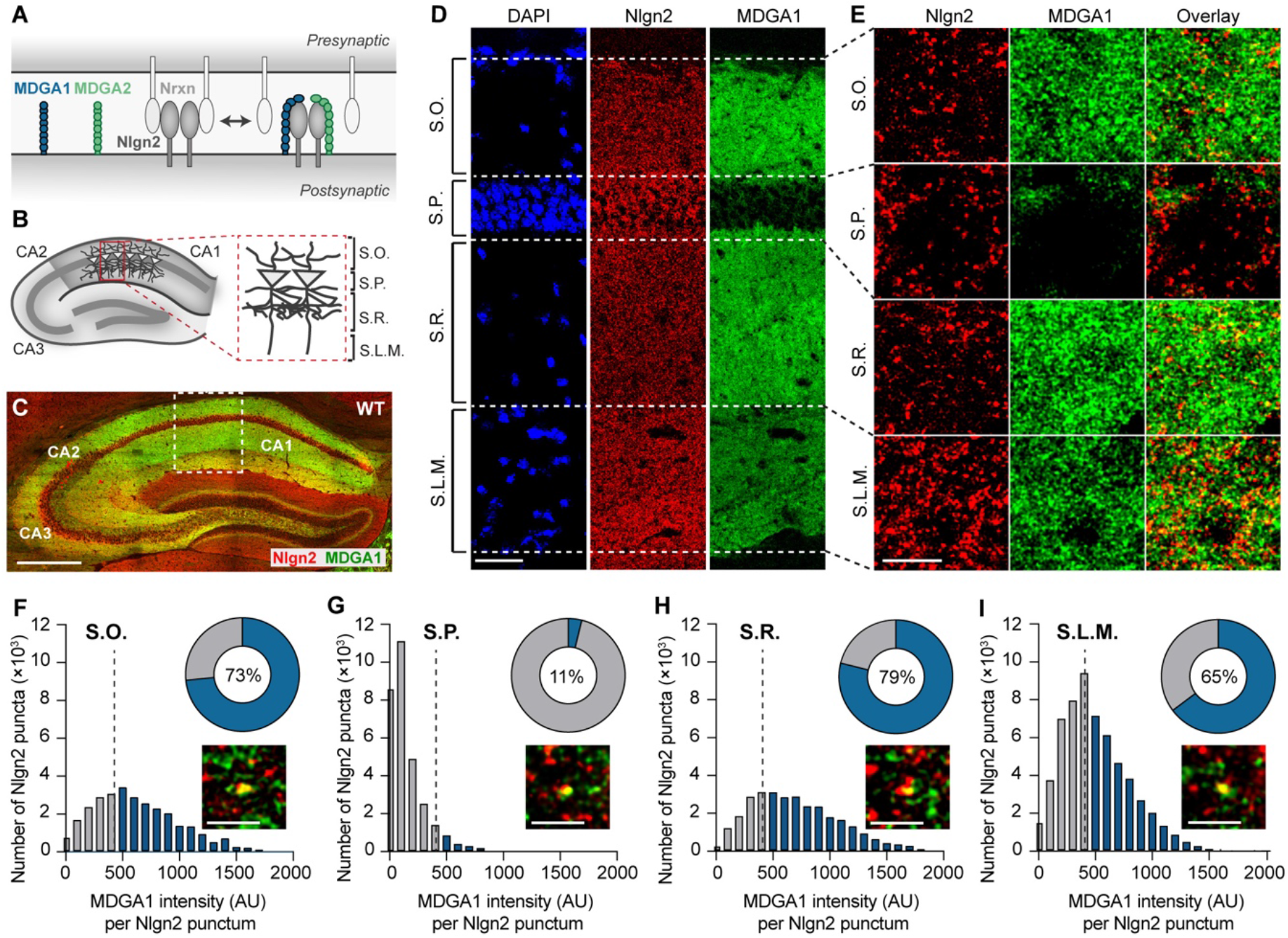
Nlgn2 and MDGA1 colocalization in hippocampal CA1 layers in WT mice. (A) Model for the putative interaction between MDGAs and Nlgn2 in the synaptic cleft. (B) Schematic representation of the dorsal hippocampus showing the layers of area CA1 in which images were acquired: Stratum oriens (S.O.), Stratum pyramidale (S.P.), Stratum radiatum (S.P.) and Stratum lacunosum-moleculare (S.L.M). (C) Photomicrograph showing a low magnification overview of the dorsal hippocampus of a WT mouse labelled with antibodies against Nlgn2 (red) and MDGA1 (green). Scale bar 500 µm. (D) Photomicrographs showing an overview of hippocampal area CA1 of a WT mouse labelled with DAPI (blue) and with antibodies against Nlgn2 (red) and MDGA1 (green). Scale bar 50 µm. (E) High magnification photomicrographs obtained from each layer showing Nlgn2 (red) and MDGA1 (green) labeling. Scale bar 5 µm. (F-I) Histograms showing the frequency distribution of MDGA1 fluorescence intensity in arbitrary units) within Nlgn2-labelled puncta in each layer. Bars in blue represent Nlgn2-labeled puncta with above-threshold MDGA1 fluorescence intensity (see Methods section for threshold determination). Doughnut chart insets display the percentage of Nlgn2-labelled puncta with an above-threshold MDGA1 fluorescence intensity (in blue, percentage in the center of the doughnut chart). High-resolution photomicrograph insets show examples of MDGA1-colocalized Nlgn2 puncta for each hippocampal layer. Scale bar 2 µm.

## Materials and methods

### Experimental subjects

Neuroligin-2 knockout (Nlgn2 KO) mice (42) were generated in our laboratory at the Max Planck Institute for Multidisciplinary Sciences (formerly Max Planck Institute of Experimental Medicine) and were maintained on a C57BL/6JRj background (Janvier Labs). MDGA1 knockout (43) and MDGA2 heterozygous knockout (25) mice on a C57BL6 background were generously provided by Tohru Yamamoto, and they were imported to the Max Planck Institute for Multidisciplinary Sciences via the laboratory of Ann Marie Craig, University of British Columbia. The mouse lines were crossed to generate Nlgn2 / MDGA1 Het or Nlgn2 / MDGA2 Het mice, and they were then backcrossed an additional 5-6 generations to a C57BL6/JRj background. For experiments involving MDGA1, Nlgn2 / MDGA1 double Het parents were crossed to generate experimental cohorts consisting of littermates of four genotypes, i.e. WT, Nlgn2 KO, MDGA1 KO and Nlgn2 / MDGA1 dKO mice. For experiments involving MDGA2, the breeding strategy needed to be adjusted, since homozygous deletion of MDGA2 is lethal (25). Therefore, one Nlgn2 Het / MDGA2 Het parent was crossed with one Nlgn2 Het / MDGA2 WT parent to generate experimental cohorts consisting of littermates of four genotypes, i.e. WT, Nlgn2 KO, MDGA2 Het and Nlgn2 KO / MDGA2 Het mice. Animals were group-housed (2-4 mice per cage) and maintained on a 12 h light/dark cycle, with food and water ad libitum, and All experiments were performed during the light cycle. Male and female mice were used for all experiments in strict sets of four sex-matched mice, one of each experimental genotype, that were strictly processed together as a set from initiation of data acquisition to completion of data analysis. For immunohistochemistry and behavior experiments, mice were 8-12 weeks old at the beginning of the experiment, while for electrophysiology experiments, mice were 6-8 weeks old. Experimenters were blind to genotype during all stage of data acquisition and analysis. All procedures were approved by the state of Niedesachsen (Landesamt für Verbraucherschutz und Lebensmittelsicherheit, license number 33.19-42502-04-18/2957) and followed the guidelines of the welfare of experimental animal use issued by the federal government of Germany and the Max Planck Society.

### Immunohistochemistry

Adult mice (8-12 weeks old) were anesthetized with isoflurane, decapitated, and their brains were rapidly dissected and immersed in isopentane at -35 to -38° C for approximately 30 sec. Brains were stored in the cryostat (Leica CM3050S, Leica Biosystems, Germany) for 30 min at -20° C, after which 14 µm thick coronal brain sections were cut and mounted on glass slides. Brain sections were arranged in experimental sets containing sex-matched mice of all four genotypes that were processed together throughout the experiment, and they were mounted such that each glass slide contained exactly one section from each of the four experimental genotypes. Sections were dried at RT for 30 min and then fixed using one of two protocols, either methanol fixation or paraformaldehyde (PFA) post-fixation, depending on the antibody used. For methanol fixation (used for anti-Nlgn2, anti-MDGA1, anti-gephyrin and anti-GABA_A_Rγ2 antibodies), sections were immersed in methanol pre-cooled to -20 C for 5 min, followed by 3x 10 min washes with PBS. For PFA post-fixation (used for the anti-VIAAT antibody), sections were incubated in 4% PFA in 0.1 M Sorensen’s phosphate buffer (pH 7.5) for 10 min at RT, followed by 2x 10 min washes with PBS and 1x 10 min wash with sodium citrate buffer (10 mM sodium citrate, 0.05% Tween-20, pH 8.0). They were then subjected to an antigen retrieval procedure, in which they were incubated in sodium citrate buffer at 95°C for 30 min, followed by a cooling period of 20 min and subsequently 2x 10 min washes with PBS at RT. After either methanol fixation or PFA post-fixation, sections were incubated for 1 hour in blocking buffer (3% bovine serum albumin, 10% goat serum and 0.3% Triton-X) at RT. Afterwards, sections were incubated overnight at 4° C in primary antibody in blocking buffer. The following primary antibodies, all from Synaptic Systems (Göttingen, Germany), were used: guinea pig anti-Nlgn2 (1:1000, Cat# 129205); rabbit anti-MDGA1 (1:2000, Cat# 421002); mouse anti-gephyrin (1:2000, Cat. # 147111); rabbit anti-GABA_A_Rγ2 (1:2000, Cat# 224003); mouse anti-VIAAT (1:2000, Cat# 131011). On the next day, sections were washed 3x 10 min with PBS and then incubated for 2 hours with the following secondary antibodies (all from ThermoFisher Scientific, USA) in blocking buffer at RT: goat anti-guinea pig, A555 (1:1200, Cat # A21435); goat anti-rabbit A488 (1:1200, Cat # A11008), goat anti-rabbit A555 (1:1200, Cat # A21429), goat anti-mouse A488 (1:1200, Cat # A11029). Sections were then washed 2x 10 min with PBS, incubated 10 min with DAPI (0.1 µg/ml, in PBS), washed 2x 10 min PBS, and stored overnight at 4° C to dry. Finally, they were coverslipped using Aqua-Poly/Mount mounting medium (Polysciences, Inc, USA).

### Image acquisition and processing for analysis of Nlgn2 and MDGA1 colocalization

Image acquisition for analysis of Nlgn2 and MDGA1 colocalization was conducted using a Leica TCS-SP8 laser scanning confocal microscope (Leica microsystems, Germany) equipped with white light laser (WLL) and hybrid detectors (HyD). A 63x oil immersion objective with a numerical aperture of 1.4 was used to obtain single plane micrographs at 1024×1024 spatial resolution and pixel spacing of xy= 45.09 nm. Laser power was optimized to ensure that the detected fluorescence intensity is within the dynamic range of detection. All imaging parameters were kept constant for images acquired from WT and Nlgn2 / MDGA1 dKO mouse brain sections and also for all images acquired from different HPC layers. Images were then subjected to deconvolution using the Lightning function of the Leica LAS X software (global mode). Tiled overview images of the hippocampus were acquired using the 20x oil immersion objective of the Leica SP8 (numerical aperture 0.75) where the navigator function of the LAS-X software was used to acquire and stitch the tiles. Tiled overview images of the CA1 were acquired using the 63x objective.

Composite two-channel images of Nlgn2 and MDGA1 labelling acquired from different hippocampus layers were split and further processed using FIJI software. Images of Nlgn2 channel were binarized and subjected to noise despeckle and watershed segmentation in FIJI to retain clearly defined Nlgn2 puncta. The threshold value used for binarization of Nlgn2 images was calculated as follows: Threshold = 20×average intensity of images acquired from the KO sample mounted on the same slide as the WT sample used in the analysis. Binary images were then subjected to segmentation using the ‘analyze particles’ algorithm of FIJI using a size filter of 0.03-1.5 µm^2^ and were added to the ROI manager where the average intensity for every Nlgn2 punctum was measured inside the MDGA1 channel by using the ‘Measure’ command while redirecting the measurement settings to the MDGA1 image. Frequency distribution histograms of MDGA1 intensity inside Nlgn2 puncta across all images acquired (8 images per layer) were generated. Total area imaged per layer was 17060.74 µm^2^. High magnification photomicrographs in Figure 1 and Supplementary Figure1 were processed by being subjected to contrast enhancement and smoothing (1x). The same minimum and maximum brightness range for every channel was used for all images taken from WT and dKO samples and for all images across hippocampus layers to allow for comparison. In order to calculate the percentage of Nlgn2 puncta colocalized with MDGA1 for every layer, a threshold of MDGA1 average fluorescence intensity was applied, defined as 10× the average intensity measured in images acquired from KO samples. The number of Nlgn2 puncta containing mean MDGA1 intensity above threshold was considered as colocalized and was then calculated as a percentage of the total number of Nlgn2 puncta detected per hippocampal layer and plotted as a doughnut chart using GraphPad Prism.

### Image acquisition and processing for quantification of gephyrin, GABA_A_Rγ2 and VIAAT puncta

Images of synaptic markers were obtained using the Leica TCS-SP8 laser scanning confocal microscope (Leica microsystems, Germany) equipped with white light laser (WLL) and hybrid detectors (HyD), 63x oil immersion objective and 4x digital zoom at spatial resolution of (512×512 pixels). Within each set of four mice, sections were anatomically matched and settings for laser power, gain and offset were kept constant during image acquisition. For each animal, 12 z-stacks, each containing 4-5 optical sections, were obtained from each layer of the dorsal hippocampus CA1 region.

For analysis of Gephyrin, GABA_A_Rγ2 and VIAAT puncta in hippocampal CA1 layers S.O., S.R. and S.L.M., images were binarized using a threshold value that was applied to all images obtained from the same experimental set of four mice. To determine the threshold value for each set, background intensity for every image was manually measured and averaged across all images belonging to the same set, and the threshold for the entire set was defined at 3x background intensity for gephyrin, GABA_A_Rγ2, and VIAAT puncta, and 10x background intensity for gephyrin aggregates. Binarized images were then subjected to noise despeckle and watershed segmentation algorithms in FIJI to reduce noise and improve segmentation to puncta. Next, images were subjected to “Analyze Particles” segmentation algorithm using a size filter of 0.04-1 µm^2^ for Gephyrin puncta, 0.8-infinity for Gephyrin aggregates and 0.04-3.25 µm^2^ for GABA_A_ R γ2 puncta. Total number, size and total intensity of puncta for every image were measured and average values were calculated per each experimental group and plotted for each hippocampal layer.

To quantify perisomatic synapses in layer S.P., the perisomatic area was manually identified by tracing the perimeter of the cell body (defined as a circular area devoid of immunofluorescence signals). The perimeter was then expanded by 1.4 µm in each direction for quantification of puncta in the perisomatic region of the outlined cell body. Synaptic puncta were quantified in the selected area using the “analyze particles” algorithm in FIJI. Number and total area of particle were normalized by the perimeter length of the cell.

### Slice electrophysiology

Young adult mice (6-8 weeks old), arranged in sex-matched sets of four genotypes, were used for experiments within a few days of each other. Mice were anesthetized with Avertin (2,2,2-Tibromoethanol, Sigma) and subsequently transcardially perfused for 100 s with ice-cold artificial cerebrospinal fluid (aCSF) containing (in mM): 124 NaCl, 2.5 KCl, 25 NaHCO_3_, 1.25 NaH_2_PO_4_) and supplemented with (in mM) 7 MgCl_2_, 0.5 CaCl_2_, 120 Sucrose (44). Mice were decapitated, and brains were rapidly dissected. Two coronal cuts were performed to isolate the hippocampal regions, which were then transferred to a chamber filled with the ice-cold sucrose-aCSF. Subsequently, tissue blocks containing the hippocampal formation were mounted and 300 μM thick coronal sections were cut on a vibratome (Leica VT1000). Slices containing the dorsal hippocampal CA1 were placed in a chamber filled with normal aCSF containing 2 mM CaCl_2_ and 1 mM MgCl_2_ (continuously bubbled with 95% O_2_ and 5% CO_2_; pH = 7.3, osmolarity = 300 mOsm). Slices were allowed to recover for 30 min at 35°C and maintained at room temperature (RT) for 4.5 hours. Chemicals were obtained from Tocris Bioscience (Bristol, Uk) and Sigma Aldrich (Darmstadt, Germany).

Whole-cell patch-clamp recordings were conducted at RT (∼22°C). During recordings, slices were continuously perfused with normal aCSF containing 2 mM CaCl_2_ and 1 mM MgCl_2_ at a rate of 1.5-2 ml/min. Hippocampal CA1 pyramidal neurons were visually identified using an upright microscope equipped with an infrared video microscopy with a 60× objective. Patch pipettes (2.5-4.0 MΩ open tip resistance) were pulled from borosilicate glass (GB150-8P Science Product, Hofheim, Germany). The holding potential was set to -65 mV. Spontaneous inhibitory synaptic currents (sIPSC) and miniature synaptic inhibitory currents (mIPSC) were recorded under voltage clamp with patch pipettes filled with an internal solution containing in (mM): 135 KCl, 15 K-gluconate, 10 EGTA, 10 HEPES, 2 MgCl_2_, 2 Na_2_-ATP (osmolarity = 332 mOsm). To pharmacologically isolate GABAergic synaptic currents, 2 μM 6-cyano-7-nitroquinoxaline-2,3-dione (NBQX, Hello Bio, Bristol, UK) and 2 μM (R)-3-(2-Carboxypiperazin-4-yl)-propyl-1-phosphonic acid ((R)-CPP, Hello Bio, Bristol, UK) were added to the bath to block AMPARs and NMDARs, respectively. During sIPSC recordings, 2 mM 4N-(2,6-Dimethylphenylcarbamoylmethyl) triethylammonium bromide (QX314, Hello Bio, Bristol, UK) was added to the internal solution to block voltage-activated Na^+^ currents. During mIPSC recordings, 1 μM tetrodotoxin (TTX, Tocris, Bristol, UK), was added to the bath solution to block voltage-activated Na^+^ currents and suppress action potential (AP) firing. Membrane resistance and cell capacitances were estimated from current transients recorded under voltage clamp in response to 10 mV depolarizing voltage steps from a holding potential of -70 mV, and calculated by assuming a simplified two-compartment equivalent circuit model (45). AP firing threshold was estimated with a ramp protocol under current clamp by injecting depolarizing current increasing from 0 pA up to 100 pA, 200 pA or 300 pA. AP phase-plane plots were constructed from the responses to the lowest depolarization exceeding AP firing threshold. During voltage clamp, a variable fraction of series resistance compensation was applied in order to maintain a residual uncompensated series resistance of 6.25 MΩ. Recordings with an initial uncompensated series resistance of >12.5 MΩ were discarded. The series resistance before compensation was allowed to change by no more than 20% during recordings. Recordings with a leak current >300 pA were discarded. The identity of visually identified CA1 pyramidal cells was confirmed based on their passive membrane properties and their discharge behaviour in response to depolarizing current steps. Patch-clamp data were acquired using an EPC-10 amplifier and Pulse or Patchmaster software (HEKA Elektronik, Germany), using a low-pass Bessel filter with at a cut-off frequency of 5 kHz and digitized at 20 kHz. All offline analyses were performed with IgorPro (Wavemetric, USA). Both sIPSC and mIPSC were detected using a sliding template-matching algorithm implemented in IgorPro (46) after additional offline filtering using a Gaussian low-pass filter with a cut-off frequency of 1 kHz.

### Behavioral analysis

Adult mice (8-12 weeks old), arranged in sex-matched sets of four genotypes that were tested on the same day, were assessed for anxiety-related behaviors in an open field test (OF) as previously described (8, 47). The OF was performed in a square arena (50 × 50 cm) made of white plastic, with a 25 × 25 cm center defined during analysis. Mice were placed in one corner and were permitted to explore the arena for 10 min. Performance was recorded using an overhead camera system and scored automatically using the Viewer software (Biobserve, St. Augustin, Germany). Between each mouse, the arena was cleaned thoroughly with 70% ethanol followed by water to eliminate any odors left by the previous mouse.

### Statistical analysis

Statistical analysis was performed using Prism (GraphPad Software, La Jolla, CA, USA) and IgorPro (Wavemetric, USA). Outliers were identified and removed using the Mean and Standard Deviation Method with a threshold value of 2, and data were subjected to two-way ANOVA with Nlgn2 genotype and MDGA genotype as the two factors. Significant main effects of Nlgn2 and/or MDGA genotype, and/or significant Nlgn2 x MDGA interactions, are reported as light grey asterisks above the corresponding graphs, and all effects are reported in Table 1 (Figures 2-5) or Supplementary Table 3 (Supplementary Figures 2-3). Post-hoc analysis was conducted using Tukey’s test for comparison between groups, and significant effects are reported as black asterisks above the corresponding comparison. * p < 0.05, ** p < 0.01, *** p < 0.001, **** p < 0.0001.

**Table 1.**
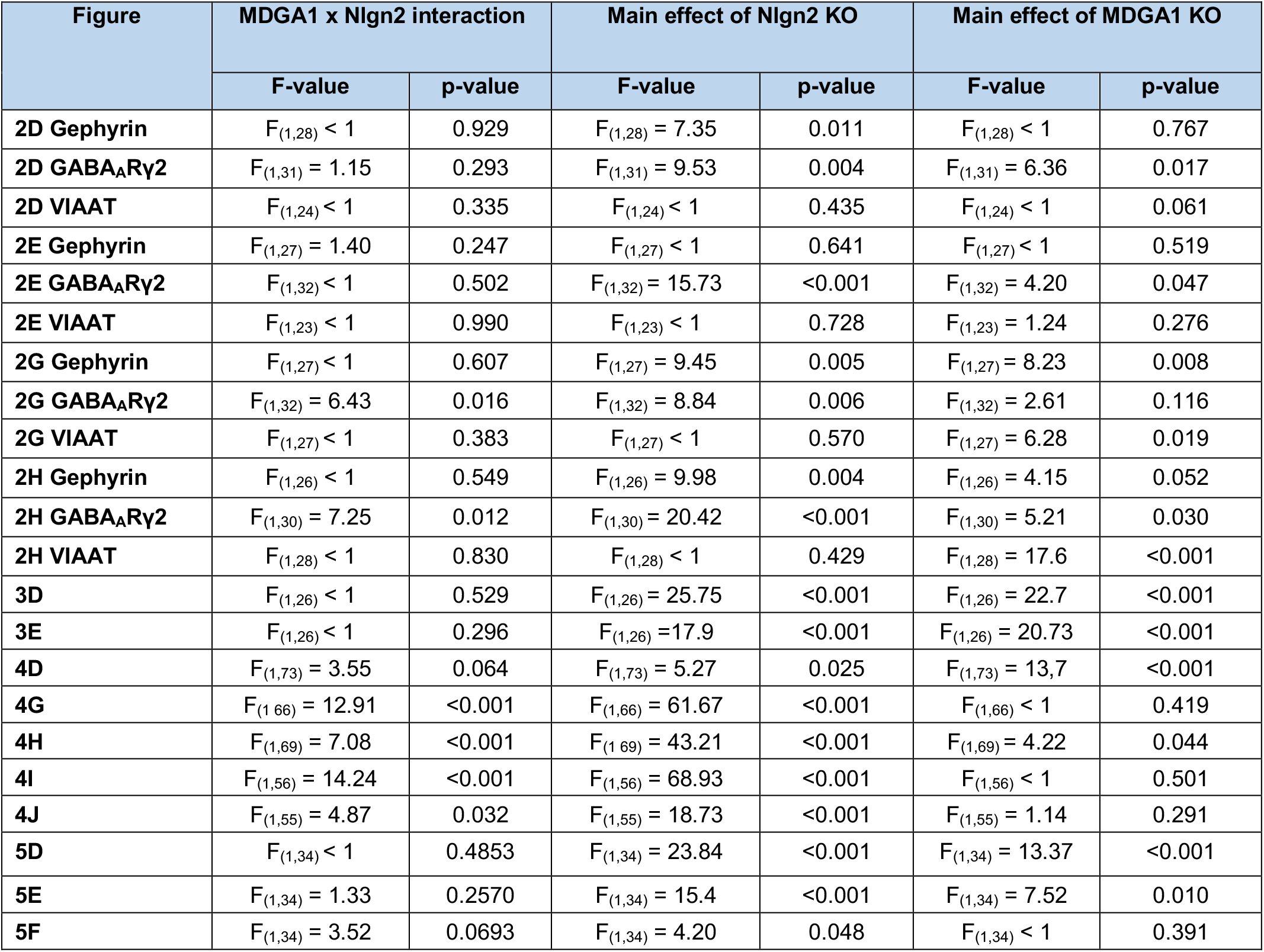
Two-way ANOVA comparisons for Figures 2-5.

**Figure 2.**
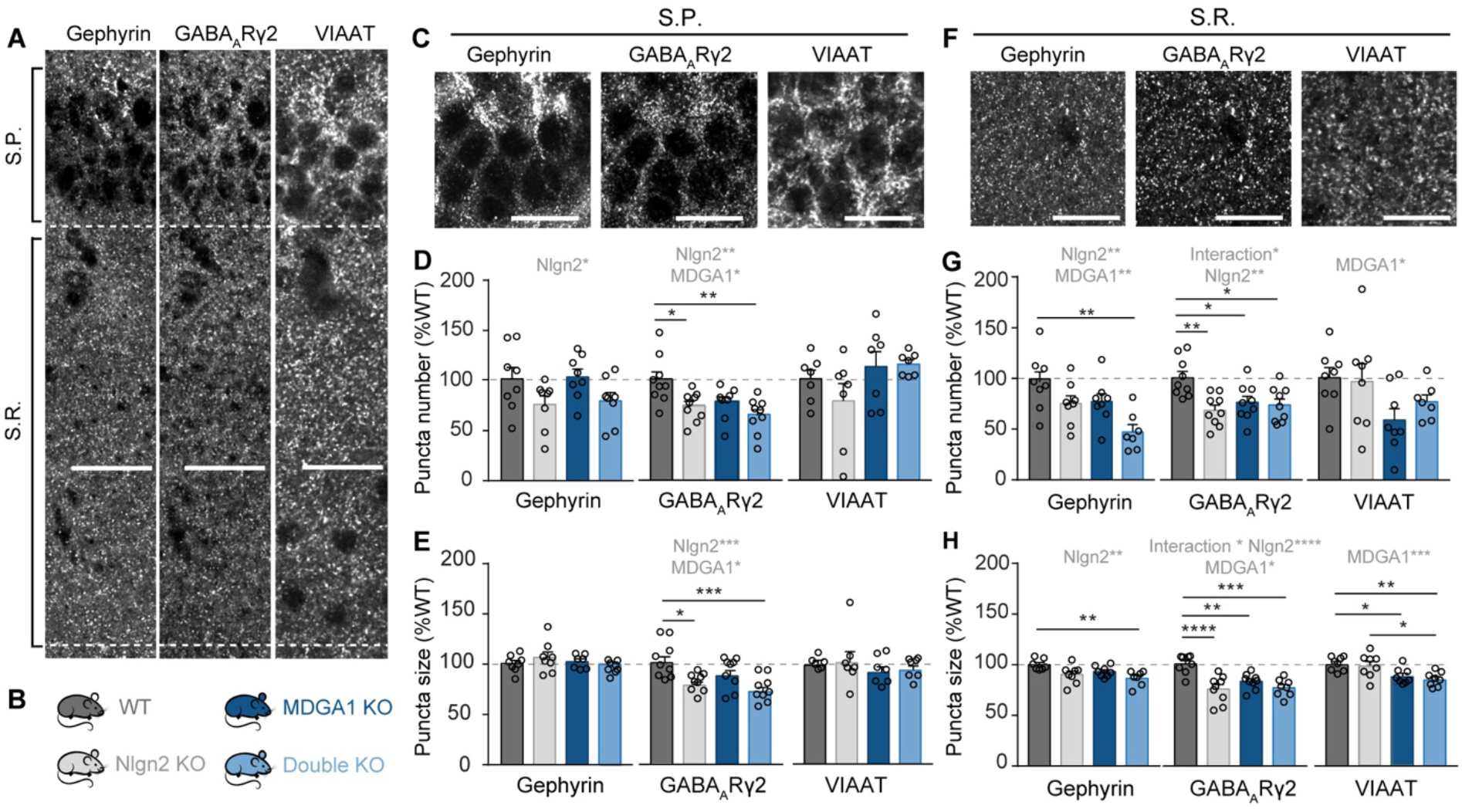
Immunohistochemical characterization of pre- and postsynaptic markers in layers S.P. and S.R. of the adult hippocampal CA1 region. (A) Photomicrographs showing an overview of layers S.P. and S.R. in a WT mouse labelled with antibodies against gephyrin (left), GABA_A_Rγ2 (middle), and VIAAT (right). Scale bar 50 µm (B) Schematic representation of the four genotypes analyzed in this study. (C) High magnification photomicrographs of layer S.P. in a WT mouse labeled with antibodies against gephyrin (left), GABA_A_Rγ2 (middle), and VIAAT (right). Scale bar 5 µm. (D-E) Quantification of the number and size of gephyrin, GABA_A_Rγ2, and VIAAT synaptic puncta in the S.P. (F) High magnification photomicrographs of layer S.R. in a WT mouse labeled with antibodies against gephyrin (left), GABA_A_Rγ2 (middle), and VIAAT (right). Scale bar 5 µm. (G-H) Quantification of the number and size of gephyrin, GABA_A_Rγ2, and VIAAT synaptic puncta in the S.R. Statistically significant ANOVA comparisons are marked in gray at the top of panels and are listed in Table 1. For all other ANOVA comparisons, F<1. Post-hoc analysis (Tukey’s comparison test): * p<0.05, ** p<0.01, *** p<0.001, **** p<0.0001. Error bars represent SEM, and each circle represents an experimental animal (n = 7-8 for gephyrin; 8-9 for GABA_A_Rγ2, 6-8 for VIAAT).

## Results

### MDGA1 co-localizes with Nlgn2 predominantly in dendritic, but not perisomatic, regions of CA1 pyramidal neurons

To determine how MDGAs may interact differentially with Nlgn2 in the regulation of GABAergic synapse function, we first used immunolabeling to assess the co-localization of MDGAs and Nlgn2 in WT mice. We focused on area CA1 in the hippocampus of adult (8-12 week old) mice (Figure 1A-B) due to the high expression level and known synaptic function of both Nlgn2 and the MDGA proteins in this region (10, 24, 25, 29, 31). Moreover, the layered structure and the precisely defined pattern of connectivity of GABAergic interneurons in area CA1 make this region uniquely suited for the study of the molecular diversity at GABAergic synapses (48). Since we were unable to identify a good antibody against MDGA2, we focused on the analysis of MDGA1 using a recently reported antibody (31). Specificity of antibodies against MDGA1 and Nlgn2 was confirmed using Nlgn2 / MDGA1 double KO (dKO) mice as a control (Supplementary Figure 1A-B). A strong expression of both MDGA1 and Nlgn2 was observed in hippocampal regions CA1, CA2 and in CA3 (Figure 1C), consistent with previous reports (31). Detailed immunolabeling analysis in CA1 revealed a prominent but relatively diffuse distribution of MDGA1 in the stratum oriens (S.O.), stratum radiatum (S.R.) and stratum lacunosum moleculare (S.L.M.), but only low levels in the stratum pyramidale (S.P.) (Figure 1D-E and Supplementary Figure 1C-F, green signal). In contrast, Nlgn2 localization was punctate and strongest in S.P. and S.L.M., and slightly weaker in S.O. and S.R. (Figure 1D-E and Supplementary Figure 1C-F, red signal). To investigate the co-localization of MDGA1 with Nlgn2 puncta in the different layers, the intensity of the MDGA1 signal within Nlgn2 puncta was calculated and plotted as a histogram showing the number of Nlgn2 puncta at each MDGA1 signal intensity (Figure 1F-I). In order to compare the number of Nlgn2 puncta that contained strong MDGA1 immunolabeling in each layer, an empirically determined threshold (signal intensity 10-fold above MDGA1 KO signal) was applied and the percentage of total Nlgn2 puncta with above-threshold MDGA1 immunolabeling was determined (Figure 1F-I, pie charts). Consistent with the expression intensity of MDGA1, the greatest degree of colocalization with Nlgn2 was observed in S.R. and S.O., less in S.L.M. and very little in S.P. Taken together, these findings indicate that MDGA1 is present primarily in dendritic but not perisomatic compartments of CA1 pyramidal neurons, and that direct molecular interactions with Nlgn2 are likely to take place in both apical and basal dendrites of these neurons, where both proteins are found.

### MDGA1 and Nlgn2 functionally interact to regulate GABAergic synapses in layer S.R. of hippocampal area CA1

While the interaction between Nlgn2 and MDGAs has been extensively investigated at the structural level (22, 23, 28), its direct functional consequences for GABAergic postsynaptic sites in intact neuronal circuits are poorly understood. To address this question, we investigated the composition of GABAergic postsynapses in hippocampal area CA1 in Nlgn2 / MDGA1 dKO mice compared to WT, Nlgn2 KO and MDGA1 KO mice (Figure 2A-B). We used immunolabeling to identify layer-specific alterations in the inhibitory synapse-specific postsynaptic scaffolding protein gephyrin and the GABA_A_R subunit γ2 (Figure 2A), which were both shown to be reduced in Nlgn2 KO mice in a synapse subtype-specific manner (10). Accordingly, we observed a reduction of gephyrin and GABA_A_Rγ2 staining in Nlgn2 KO mice which covered layer S.P. (Figure 2C-E, grey bars, and Table 1), but surprisingly extended to layer S.R. (Figure 2F-H, grey bars, and Table 1) and, to a lesser degree, to layer S.O. (Supplementary Table 1). In contrast, MDGA1 KO affected gephyrin and GABA_A_Rγ2 staining most prominently in layer S.R. (Figure 2F-H, dark blue bars, and Table 1), in keeping with our observation that MDGA1 is localized most strongly in this layer (Figure 1D-E). In particular, MDGA1 KO mice displayed a trend toward a reduction in the number and size of gephyrin puncta specifically in layer S.R. (Figure 2F-H, dark blue bars, and Table 1), and this trend was strongly exacerbated in the Nlgn2 / MDGA1 dKO mice (Figure 2F-H, light blue bars, and Table 1). No effect of the MDGA1 KO or its interaction with the Nlgn2 KO was observed in any other layer (Figure 2C-E and Supplementary Table 1). Similarly, MDGA1 KO most strongly, albeit not exclusively, affected GABA_A_Rγ2 puncta in layer S.R. (Figure 2F-H, dark blue bars, and Table 1). A reduction in both the number and size of GABA_A_Rγ2 puncta was observed in single MDGA1 KO mice, which matched the reduction observed in Nlgn2 KO mice and was not further exacerbated in the Nlgn2 / MDGA1 dKO mice. More subtle effects of MDGA1 deletion were observed in layers S.P. and S.L.M., with a reduction of the number and size of GABA_A_Rγ2 puncta in layer S.P. and of the number of GABA_A_Rγ2 puncta in layer S.L.M. (Figure 2C-E, dark blue bars, and Supplementary Table 1, respectively). Together, these findings highlight that MDGA1 deletion most prominently affects GABAergic postsynapses in layer S.R. of hippocampal area CA1, and that interactions between the effects of MDGA1 and Nlgn2 are restricted to this layer.

To clarify whether Nlgn2 and/or MDGA1 selectively regulate the composition of postsynaptic sites, or whether they also play a role at presynaptic terminals, we performed immunohistochemical analysis for the presynaptic vesicular inhibitory amino acid transporter (VIAAT). Consistent with previous reports (8, 10), deletion of Nlgn2 did not alter size or number of VIAAT puncta (Figure 2C-H, grey bars, and Supplementary Table 1). In contrast, deletion of MDGA1 resulted in a small but significant decrease in the size of VIAAT puncta, most prominently in layers S.R., and S.O., to a lesser extent in layer S.L.M, but not in layer S.P. (Figure 2C-H, dark blue bars, and Supplementary Table 1). This pattern is highly consistent with the differential expression of MDGA1 in these hippocampal layers. Together, our findings indicate that Nlgn2 and MDGA1 mediate largely distinct and layer-specific effects at GABAergic synapses in hippocampal area CA1, which reflect their differential expression in the respective layers. Interactions between Nlgn2 and MDGA1 function at postsynaptic sites are limited to layer S.R., where the highest degree of colocalization of Nlgn2 with MDGA1 is observed.

### MDGA2 and Nlgn2 functionally interact to regulate GABA_A_R abundance in area CA1

While most evidence indicates that MDGA1, but not MDGA2, plays an important role at hippocampal GABAergic synapses (24, 25), MDGA2 was also reported to be present at GABAergic synapses in dissociated neuron cultures (30). Unfortunately, the lack of a suitable antibody prevented us from assessing Nlgn2-MDGA2 colocalization *in situ*. To nevertheless determine whether MDGA2 modulates Nlgn2 functions at GABAergic synapses in hippocampal area CA1, we immunohistochemically assessed gephyrin, GABA_A_Rγ2 and VIAAT in Nlgn2 KO / MDGA2 heterozygous KO mice (since homozygous MDGA2 KO is lethal (25)). Intriguingly, MDGA2 Het mice displayed a reduction of the number and size of GABA_A_Rγ2 puncta that was most prominent in the S.P., and that was exacerbated in Nlgn2 KO / MDGA2 Het mice (Supplementary Table 2). No relevant effects on gephyrin or VIAAT were observed. Although it is difficult to interpret the reduction in GABA_A_Rγ2 puncta without knowing in which layers MDGA2 is expressed, it is conceivable that MDGA2 functionally replaces MDGA1 in layer S.P., from which MDGA1 is mostly absent.

**Table 2.** Passive and AP properties of CA1 pyramidal cells in WT, Nlgn2 KO, MDGA1 KO and Nlgn2 / MDGA1 dKO mice.

|  | WT |  | Nlgn2 KO |  | MDGA1 KO |  | Nlgn2 / MDGA1 dKO |  | Main source of variation |  |
| --- | --- | --- | --- | --- | --- | --- | --- | --- | --- | --- |
| | n | Mean ± SEM | n | Mean ± SEM | n | Mean ± SEM | n | Mean ± SEM | F-value | p-value |
| **Membrane resistance (MOhm)** | 37 | 100.50 ± 4.50 | 34 | 95.02 ± 5.64 | 41 | 114.608 ± 7.43 | 36 | 100.897 ± 7.65 | \ | \ |
| **Membrane capacitance, proximal compartments (pF)** | 37 | 42.89 ± 2.39 | 34 | 45.06 ± 2.66 | 41 | 39.46 ± 1.64 | 36 | 46.88 ± 1.82 | Nlgn2 KO:<br>$F_{(1,144)} = 5.05$ | 0.026 |
| **Membrane capacitance, distal compartments (pF)** | 37 | 122.74 ± 4.84 | 34 | 113.24 ± 6.03 | 41 | 111.19 ± 3.39 | 36 | 126.60 ± 5.12 | Interaction:<br>$F_{(1,144)} = 6.64$ | 0.011 |
| **Resting membrane potential (mV)** | 21 | -58.30 ± 1.72 | 16 | -57.21 ± 2.21 | 23 | -55.37 ± 1.26 | 18 | -59.92 ± 1.48 | \ | \ |
| **AP threshold (mV)** | 20 | -44.38 ± 0.77 | 15 | -43.65 ± 0.73 | 24 | -45.40 ± 0.67 | 18 | -44.65 ± 0.72 | \ | \ |
| **AP amplitude (mV)** | 20 | 117.91 ± 1.21 | 15 | 117.81 ± 1.48 | 24 | 120.863 ± 1.83 | 19 | 120.87 ± 1.40 | \ | \ |

### MDGA1 but not MDGA2 regulates the formation of Nlgn2 KO-related extrasynaptically-located gephyrin aggregates

Beyond the effects of Nlgn2 KO on GABAergic synapses, a striking and robust observation in Nlgn2 KO mice is the presence of prominent cytoplasmic gephyrin aggregates that are found at the border between layers S.P. and S.O. (10). These aggregates were proposed to result from a loss of nucleation sites for GABAergic postsynapses in the absence of Nlgn2, leading to disrupted gephyrin transport to synaptic sites (10), but whether other molecules are involved in the regulation of cytoplasmic gephyrin aggregates remains to be determined.

To investigate an involvement of MDGAs in this process, we tested whether MDGA1 or MDGA2 alone engage in the formation of these gephyrin aggregates, and whether they influence aggregate formation in Nlgn2 KO mice (Figure 3A-B). As expected based on previous reports (10), a robust increase in the number and in the total area of extrasynaptic cytoplasmic gephyrin aggregates was observed in putative CA1 pyramidal cell dendrites at the border between layers S.P. and S.O. in Nlgn2 KO mice (Figure 3C-F, grey bars, and Table 1). Strikingly, this increase was completely absent in Nlgn2 / MDGA1 dKO mice, which showed exactly the same number and total area of aggregates as WT mice (Figure 3C-F, light blue bars, and Table 1), indicating that deletion of MDGA1 reverses the effect of Nlgn2 deletion on gephyrin aggregation. In contrast, MDGA2 heterozygous deletion had no effect on gephyrin aggregation, and Nlgn2 KO / MDGA2 Het mice showed the same number and total area of aggregates as MDGA2 single KO mice (Supplementary Figure 2A-C, and Supplementary Table 3). Together, these observations indicate that MDGA1, but not MDGA2, is actively involved in retaining gephyrin in cytoplasmic aggregates through an as yet unknown mechanism.

**Figure 3.**
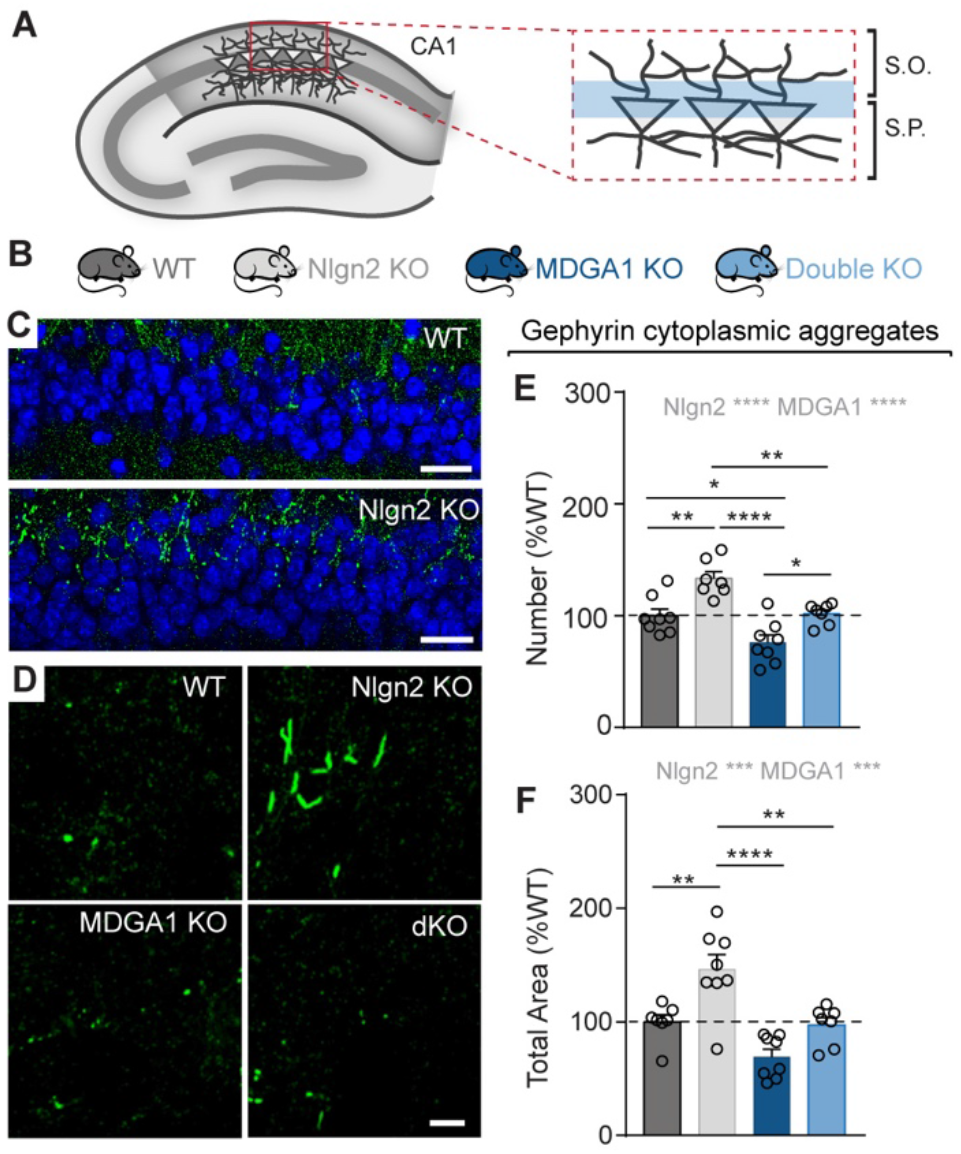
MDGA1 regulates the formation of Nlgn2 KO-related cytoplasmic gephyrin aggregates. (A) Schematic diagram of the dorsal hippocampus showing the region at the border between S.P. and S.O. in which gephyrin aggregates occur. (B) Schematic representation of the four genotypes analyzed in this study. (C) Photomicrographs of the border region between S.P. and S.O. labelled with DAPI (blue) and anti-gephyrin antibody (green) in WT and Nlgn2 KO mice. Scale bars 25 µm. (D) High magnification photomicrographs of gephyrin aggregates in WT, Nlgn2 KO, MDGA1 KO and Nlgn2 / MDGA1 dKO mice. Scale bar 5 µm. (E-F) Quantification of the number and total area of gephyrin aggregates, expressed as percentage of WT. Statistically significant ANOVA comparisons are marked in gray at the top of panels and listed in Table 1. For all other ANOVA comparisons, F<1. Post-hoc analysis (Tukey’s comparison test): * p<0.05, ** p<0.01, *** p<0.001, **** p<0.0001. Error bars represent SEM, and each circle represents an experimental animal (n = 7-8).

### Loss of MDGA1 expression perturbs GABAergic synaptic transmission in CA1 pyramidal neurons

To determine whether the observed alterations in gephyrin and GABA_A_Rγ2 puncta affect inhibitory synaptic transmission in hippocampal area CA1, we performed whole-cell patch-clamp recordings from pyramidal neurons in acute hippocampal slices obtained from adult (6-8 week old) mice (Figure 4A).

**Figure 4.**
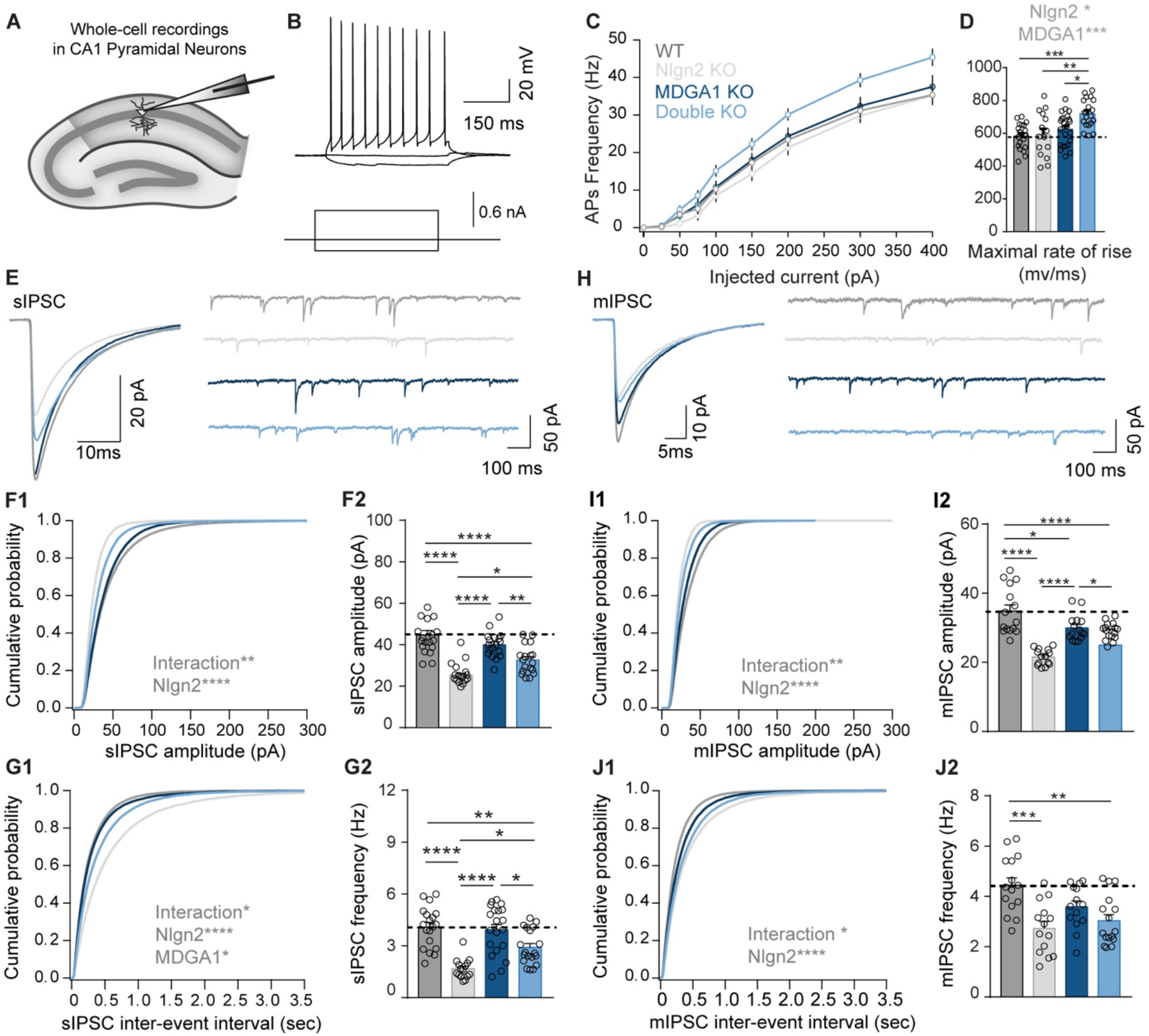
Loss of MDGA1 expression perturbs GABAergic transmission in CA1 Pyramidal neurons. (A) Schematic diagram showing location of sIPSC and mIPSC recordings in area CA1 of the adult dorsal hippocampus. (B) Representative firing pattern of CA1 pyramidal neurons. (C) Frequency of action potentials (APs) in response to steps of injected current. (D) Quantification of the maximal rate of AP rise in WT, Nlgn2 KO, MDGA1 KO, Nlgn2 / MDGA1 dKO mice. (E) Representative average sIPSCs and sIPSC recording traces from all four genotypes analysed. (F-G) Average cumulative distribution and bar-graphs showing the quantification of sIPSC amplitude (F) and sIPSC inter-event frequency (G). (H) Representative average mIPSCs and mIPSC recording traces from all four genotypes analysed. (I-J) Average cumulative distribution and bar-graphs showing the quantification of mIPSC amplitude (I) and mIPSC inter-event frequency (J). Error bars represent SEM. Statistically significant ANOVA comparisons are marked in gray at the top of panels and listed in Table 1. For all other ANOVA comparisons, F<1. Post-hoc analysis (Tukey’s comparison test): * p<0.05, ** p<0.01, *** p<0.001, **** p<0.0001. In bar graphs, each circle represents a single cell (n = 14-24 cells for APs and rate of rise; 15-20 cells for sIPSC recordings; 14-16 cells for mIPSC recordings; each from four animals per genotype).

CA1 pyramidal neurons were identified based on their location, morphology and firing pattern (Figure 4B). Their resting membrane potential and passive membrane properties were similar in all genotypes (Table 2). While action potential (AP) firing threshold, AP amplitude, and AP half width were unchanged (Table 2), CA1 pyramidal cells in Nlgn2 / MDGA1 dKO mice showed a surprising tendency towards higher AP frequency in response to injection of depolarizing current steps as compared to other genotypes (Figure 4C, light blue trace, and Table 1). Analysis of AP kinetics revealed an increased maximal rate of rise in CA1 pyramidal cells of Nlgn2 / MDGA1 dKO mice (Figure 4D, light blue bars, and Table 1). No differences in passive membrane properties and discharge behavior were detected in CA1 pyramidal cells upon MDGA2 KO (Supplementary Figure 3D-F, and Supplementary Table 4).

In line with previous observations (10), the frequency and amplitude of spontaneously occurring inhibitory synaptic events (sIPSCs; recorded in the absence of TTX) and miniature inhibitory synaptic events (mIPSCs, recorded in the presence of TTX) were strongly reduced in CA1 pyramidal cells of Nlgn2 KO mice (Figure 4E-J, grey traces, and Table 1). Surprisingly, we also observed a significant reduction of mIPSC amplitude and frequency in CA1 pyramidal cells of MDGA1 KO mice (Figure 4H-J, dark blue traces and bars, and Table 1), in contrast to previous studies reporting augmentation GABAergic inhibition following MDGA1 KO (24). Importantly, sIPSC frequency and amplitude were significantly increased in CA1 pyramidal cells of Nlgn2 / MDGA1 dKO mice compared to Nlgn2 KO mice, indicating that additional MDGA1 KO partially rescues the effects of Nlgn2 deletion on GABAergic synaptic transmission (Figure 4E-G, light blue traces and bars, and Table 1). A rescue of GABAergic transmission in Nlgn2 / MDGA1 dKO mice was also observed with respect to mIPSC frequency and amplitude, albeit to a lesser degree (Figure 4I-J, light blue, and Table 1). In contrast, loss of a single MDGA2 allele had no major effects on GABAergic transmission (Supplementary Figure 2 F-K, dark green bars, and Supplementary Table 3), nor did it rescue the functional deficits observed in Nlgn2 KO mice (Supplementary Figure 2 F-K, light green, and Supplementary Table 3).

Together, these findings indicate that single KO of MDGA1, but not of MDGA2, slightly reduces GABAergic synaptic transmission, while combined loss of MDGA1 and Nlgn2 partially reverses the profound defects of GABAergic synaptic transmission observed after Nlgn2 KO.

### MDGA1 deletion ameliorates abnormal anxiety-related avoidance behavior in female Nlgn2 KO mice

Finally, in light of the role of both Nlgn2 and MDGA1/2 in the pathophysiology of psychiatric disorders (32-41), we assessed how the interaction between Nlgn2 and MDGAs might influence psychiatrically relevant behaviors. Nlgn2 KO causes a profound increase in anxiety-related avoidance behaviors in mice (8, 47, 49), which at least partially originates from altered connectivity in hippocampal-amygdala-prefrontal circuits (50). Using an open field task, we tested whether this anxiety phenotype is modulated in Nlgn2 / MDGA1 dKO or Nlgn2 KO / MDGA2 Het mice (Figure 5 A-C, Supplementary Figure 3 A-C). Strikingly, but consistent with the amelioration of the defects in gephyrin aggregation and sIPSC frequency and amplitude, female Nlgn2 / MDGA1 dKO mice showed an amelioration of the profound Nlgn2 KO anxiety phenotype as indicated by a normalization of the time spent and the distance traveled in the center of the open field chamber (Figure 5D, F, light blue bars vs. grey bars, and Table 1). Surprisingly, however, male Nlgn2 KO mice in this experiment did not display the anxiety phenotype previously observed, likely due to complex interactions with strain background or parental behavior of the Nlgn2 Het / MDGA1 Het breeders (Supplementary Figure 3D-E and Supplementary Table 3). Accordingly, we were unable to determine whether the anxiety phenotype of Nlgn2 KO mice is also ameliorated in male Nlgn2 / MDGA1 dKO mice. No amelioration of the Nlgn2 KO anxiety phenotype was observed in male or female Nlgn2 KO / MDGA2 Het mice (Supplementary Figure 3F-H and Supplementary Table 3), consistent with the lack of an effect of MDGA2 Het on gephyrin aggregation and GABAergic synaptic transmission (Supplementary Figure 2A-C and Supplementary Table 3).

**Figure 5.**
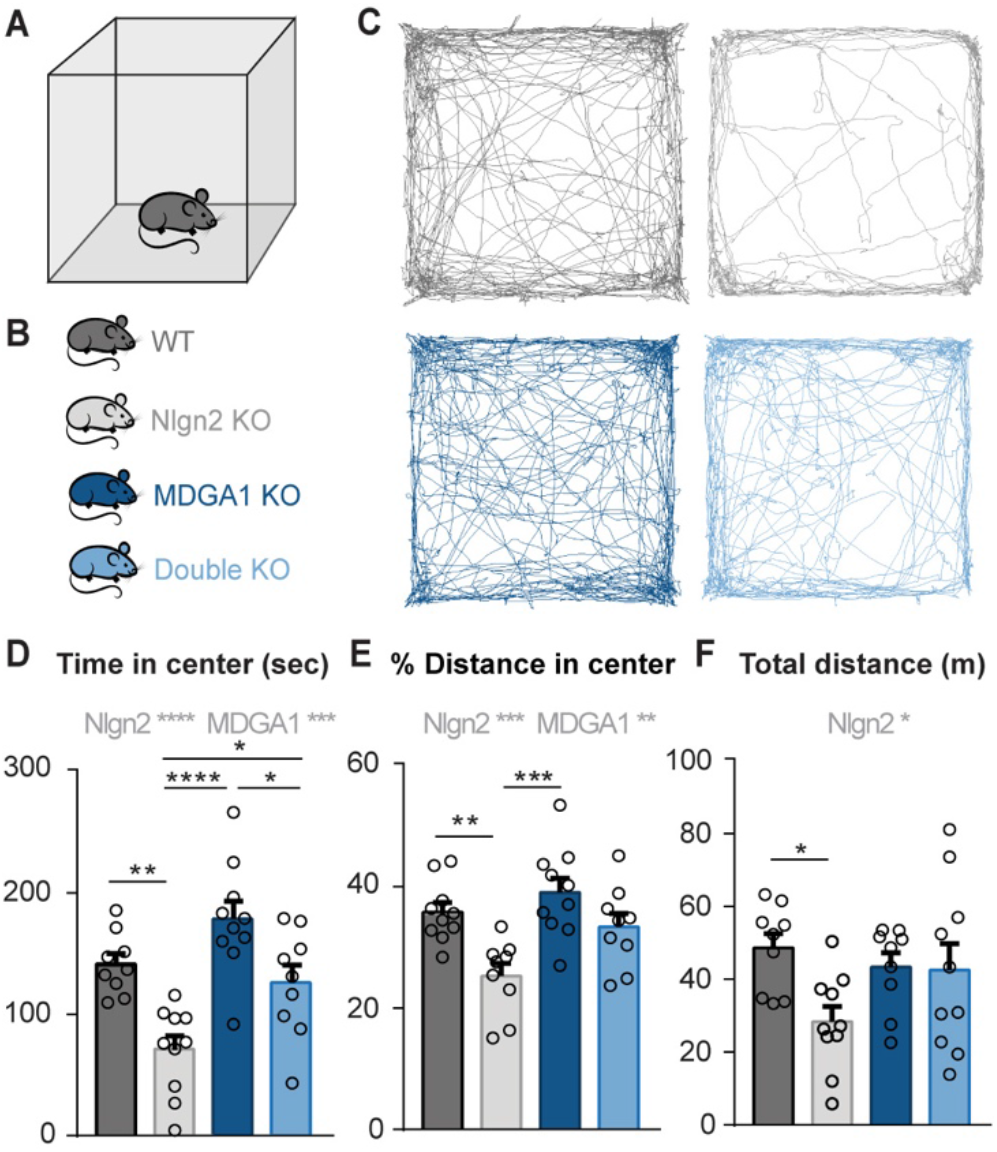
MDGA1 deletion ameliorates abnormal anxiety-related avoidance behavior in female Nlgn2 KO mice. (A-B) Schematic representation of the OF arena (A) and of the four experimental genotypes analyzed (B). (C) Representative tracks of OF exploration. (D) Time spent in the anxiogenic region (center) of the OF arena. (E) Distance traveled in the center of the OF, expressed as percentage of total distance traveled. (F) Total distance travelled in the OF. Statistically significant ANOVA comparisons are marked in gray at the top of panels and listed in Table 1. For all other ANOVA comparisons, F<1. Post-hoc analysis (Tukey’s comparison test): * p<0.05, ** p<0.01, *** p<0.001, **** p<0.0001. Error bars represent SEM, and each circle represents an experimental animal (n = 9-10).

Taken together, our observations indicate functional interactions between MDGA1 and Nlgn2 KOs, but not between MDGA2 and Nlgn2 KOs, in modulating the neuronal circuit that regulate anxiety-related avoidance behaviors. Further, our data reveal an anxiolytic effect of MDGA1 deletion in Nlgn2 KO mice, raising the intriguing possibility that the MDGA1-Nlgn2 interaction may serve as a novel target for anxiolytic therapeutic strategies.

## Discussion

In the present study, we sought to determine how the schizophrenia- and autism-associated synaptic adhesion proteins Nlgn2 and MDGAs functionally interact *in vivo* to regulate GABAergic synapses. We report that Nlgn2 and MDGA1 show distinct distribution patterns in hippocampal area CA1, with Nlgn2 localized in a punctate pattern that was most prominent in layers S.P. and S.L.M, and MDGA1 distributed diffusely throughout this region, most strongly in layers S.R. and S.O. Using immunolabeling, electrophysiological and behavioral analysis in Nlgn2 / MDGA1 dKO mice, we found that the combined loss of Nlgn2 and MDGA1 leads to an exacerbated reduction of gephyrin puncta specifically in layer S.R. At the same time, a normalization of cytoplasmic gephyrin aggregates at the S.P.-S.O. boundary was observed in Nlgn2 / MDGA1 dKO mice, accompanied by a partial normalization of defects in sIPSC characteristics in CA1 pyramidal neurons and of the anxiety-related behavior in an open field test. Heterozygous KO of MDGA2 in Nlgn2 KO mice had only very subtle effects on GABAergic synapses, indicating that MDGA2 plays a minor role in modulating Nlgn2 function. Together, our experimental data indicate that the key function of the interaction between Nlgn2 and MDGA1 bidirectionally regulates gephyrin aggregation in the intact hippocampal area CA1 network, which in turn determines the effects of these proteins on GABAergic synapse assembly and function and on anxiety-related behavior.

While extensive structural data document a molecular interaction between Nlgn2 and MDGAs *in vitro*, it has to date remained unknown to which extent and where these proteins colocalize *in situ*. Here we report that in hippocampal area CA1 of adult mice, MDGA1 shows a relatively diffuse staining pattern, which is most prominent in layers S.R. and S.O., and to a lesser extent in layer S.L.M., while it is largely absent from layer S.P. In contrast, Nlgn2 is present in a punctate pattern, consistent with previous reports (10), with a particularly prominent staining in layers S.P. and S.L.M. and a lesser, albeit still strong, staining in layers S.O. and S.R. Accordingly, the strongest colocalization of Nlgn2 with MDGA1 is observed in layer S.R., as well as in layers S.O. and S.L.M., while very little colocalization was observed in layer S.P. Consistent with their expression, the most pronounced effects of MDGA1 KO and Nlgn2 / MDGA1 dKO were observed in the S.R., with a significant reduction of GABA_A_Rγ2 in both MDGA1 KO and Nlgn2 / MDGA1 dKO mice, and an exacerbation of gephyrin loss in the Nlgn2 / MDGA1 dKO. Whether this potentiation of the loss of gephyrin clusters in Nlgn2 / MDGA1 dKO mice results from a direct molecular interaction of the two proteins, or whether it reflects independent regulatory pathways, cannot easily be distinguished. Nevertheless, our findings identify layer S.R. as one of the primary sites of the combined function of Nlgn2 and MDGA1 at GABAergic synapses in area CA1. Layer S.R. consists primarily of dendritic arborizations of pyramidal neurons with a cell body in layer S.P., and this layer is the primary area of excitatory input from area CA3 through the Schaffer collateral pathway, placing the Nlgn2-MDGA1 interaction in an ideal position to modulate the flow of information through this pathway. A number of different GABAergic neuron subtypes target the dendritic tree of CA1 pyramidal neurons specifically in layer S.R., including parvalbumin-positive bistratified cells, neuropeptide Y-positive Ivy cells, and cholecystokinin-positive Schaffer collateral-associated and apical dendrite-innervating cells (48). Whether Nlgn2 and/or MDGA1 loss selectively or predominantly affects any of these inputs remains to be determined.

A second striking effect of the Nlgn2 / MDGA1 dKO is the normalization of the number of the cytoplasmic gephyrin aggregates that occur upon Nlgn2 KO at the boundary between layers S.P. and S.O. (10). These extrasynaptic cytoplasmic gephyrin aggregates are thought to result from the loss of the gephyrin-collybistin-Nlgn2 triad, which is necessary for Nlgn2-mediated recruitment of gephyrin to GABAergic synapses and the consequent somatic gephyrin accumulation (10). However, it is unknown whether these aggregates consist exclusively of gephyrin, and why they are localized so specifically at the boundary between layers S.P. and S.O. rather than being distributed throughout the cytoplasm of CA1 pyramidal cells. Moreover, our data from Nlgn2 / MDGA1 dKO mice indicate that the presence of these cytoplasmic gephyrin aggregates appears to be unrelated to the localization of gephyrin and GABA_A_Rγ2 at synapses, since the number of gephyrin aggregates is normalized in these mice (Figure 3C-E), while the loss of synaptic gephyrin and GABA_A_Rγ2 clusters in S.R. is exacerbated (Figure 2F-H). The normalization of the cytoplasmic gephyrin aggregates correlates well with the partial normalization of sIPSC frequency and amplitude, as well as with the partial normalization of the anxiety behavior in the female Nlgn2 x MDGA1 dKO mice. These data lead to the fascinating conclusion that it is this reduction of gephyrin aggregates, rather than the exacerbation of the loss of gephyrin from layer S.R., that dominates the functional consequences of combined deletion of Nlgn2 and MDGA1. While the mechanistic link between the cytoplasmic gephyrin aggregates and the functional consequences at the cellular and behavioral level remains to be established, regional differences in the expression of gephyrin splice forms (51) or the differential recruitment and trafficking of GABA_A_R subunits other than GABA_A_Rγ2 may play a role.

Our findings raise the intriguing question as to how MDGA1 differentially regulates the number of synaptic gephyrin and GABA_A_Rγ2 clusters vs. the cytoplasmic gephyrin aggregates. Co-staining of Nlgn2 and MDGA1 indicates that MDGA1 is not strictly localized to Nlgn2-positive clusters, but rather displays a relatively diffuse distribution that partially overlaps with Nlgn2. This is consistent with recent findings in neuronal cultures indicating that MDGAs are homogeneously distributed over the cell surface and exhibit fast diffusion throughout the dendritic membrane, where they interact with Nlgn1 extrasynaptically and prevent Nlgn1 and AMPARs from entering nascent glutamatergic synapses (31). Based on our immunolabeling analysis, it is plausible that a similar mechanism holds true for the Nlgn2-MDGA1 interaction, and that direct molecular interactions primarily take place extrasynaptically to regulate the trafficking of GABAergic synapse components to synaptic sites. At the same time, it was recently shown that Nlgn2 is not the only target of MDGA1, and that a transsynaptic interaction between MDGA1 and APP regulates synapse function independently of Nlgn2 at GABAergic synapses formed by oriens-lacunosum moleculare (O-LM) interneurons onto the distal dendrites of pyramidal neurons in layer S.L.M. (29). It is conceivable that the additive effects of Nlgn2 / MDGA1 dKO in layer S.R. results from the loss of an MDGA1 interaction with an unidentified additional synaptic partner, independently of Nlgn2. In contrast, the normalization of gephyrin aggregation in the Nlgn2 / MDGA1 dKO mice likely results from the loss of a direct antagonistic interaction between these two proteins. Since MDGA1 can also bind to Nlgn1 (31) and potentially other Nlgns, it is possible that additional KO of MDGA1 in the Nlgn2 KO mice releases an excess pool of another Nlgn isoform, which can then substitute for the absent Nlgn2 in disassembling gephyrin aggregates.

A further aim of our study was to discern whether MDGA1 and MDGA2 differ in their relative importance for the modulation of Nlgn2 function at GABAergic synapses, since the relative effect of the two MDGAs on GABAergic synapse function has been controversially discussed. One set of studies on the hippocampal area CA1 indicated that deletion of MDGA1 causes an increase in mIPSC frequency but not in mEPSC frequency, and an increase in the number of symmetric but not asymmetric synapses identified in electronmicrographs (24), while heterozygous deletion of MDGA2 causes morphological and functional alterations at glutamatergic but not GABAergic synapses (25). These data led to the conclusion that MDGA1 and MDGA2 must be specific to inhibitory and excitatory synapses, respectively. In contrast, a proteomics study on cortical neuron cultures detected MDGA1 and MDGA2 in the excitatory and inhibitory synaptic clefts, respectively (30), resulting in the opposite conclusion. Recent data from hippocampal cultures indicated no change in mIPSC frequency or amplitude following deletion of either MDGA1 or MDGA2 (31), leading the authors to conclude that during early development, neither MDGA protein plays a role at inhibitory synapses. A further recent study in hippocampal area CA1 indicated that overexpression of WT MDGA1 and Nlgn2 binding-deficient MDGA1, but not APP binding-deficient MDGA1, causes a reduction in mIPSC frequency, while conditional deletion of MDGA1 in area CA1 had no effect on mIPSC frequency (29). These findings indicate that the effects of the MDGAs appear to depend on the experimental conditions, potentially due to differences in the ratio of MDGAs to Nlgns and other binding partners as shown previously (31). Here we report subtle and layer-specific effects on GABA_A_Rγ2 localization in both MDGA1 KO and MDGA2 Het mice, but the most prominent effects on gephyrin aggregates, sIPSC and mIPSC frequency, were found almost exclusively in Nlgn2 / MDGA1 KO but not Nlgn2 KO / MDGA2 Het mice. Surprisingly, deletion of only MDGA1 resulted in a modest reduction in mIPSC amplitude as well as in a decreased size of VIAAT and GABA_A_Rγ2 puncta size, in stark contrast to the increase in mIPSC frequency and symmetric synapse density previously reported in area CA1 (24). The reason for this discrepancy most likely lies in differences in the strain background (C57BL/6JRj in our study, mixed C57BL/6NxJ in Connor et al. (24)), which has previously been shown to modulate both MDGA2 function (52) and GABA_A_Rα2 expression (53). Importantly, our findings of a reduction of GABAergic synapse markers and smaller mIPSC amplitude are internally consistent, and our observation of a prominent reduction of sIPSC and mIPSC frequency and amplitude, as well as gephyrin and GABA_A_Rγ2 staining intensity in the Nlgn2 KO, are consistent with previous findings (10), validating our technical approach. Together, our data indicate that in adult hippocampal area CA1, MDGA1, and not MDGA2, interacts with Nlgn2 to regulate GABAergic synapses, and that under our conditions, MDGA1 deletion alone results in a decrease, rather than an increase, in GABAergic synapse function in area CA1.

The ultimate objective of our study has been to identify the key mechanisms by which Nlgn2 and MDGAs contribute to psychiatrically relevant phenotypes of the psychiatric disorders they have been linked to, such as autism and schizophrenia (7, 20, 32-41), and to determine how such mechanisms can be targeted to ameliorate pathophysiological behaviors. We show that combined deletion of Nlgn2 and MDGA1 results in a partial reversal of the prominent anxiety phenotype observed in female Nlgn2 KO mice, consistent with the normalization of gephyrin aggregates and the partial normalization of sIPSC frequency in CA1 pyramidal neurons. Given this correlation, it is conceivable that the formation of these gephyrin aggregates is at the core of the anxiety phenotype in the Nlgn2 KO mice, and that it may be possible to reverse the pathophysiological consequences of Nlgn2 or MDGA1 variants by reversing gephyrin aggregate formation. A further detailed understanding of how Nlgn2 and MDGA1 differentially regulate gephyrin aggregation, and how these aggregates affect GABAergic synapse function, will therefore be critical towards developing new therapeutic approaches for Nlgn2- and MDGA1-related psychiatric disorders.

## Acknowledgments

This study was supported by the Deutsche Forschungsgemeinschaft (Heisenberg Grant, D.K-B.; EXC2067/1-390729940, N.B.), the Volkswagen Foundation (Experiment! Grant, D.K-B.), and the German Federal Ministry of Education and Research (ERA-NET Neuron Synpathy, N.B.). T.Z. was a student at the Mainz Research School of Translational Biomedicine (TransMed), funded by a Ph.D. fellowship from the Focus Program Translational Neurosciences (FTN). H.A was a student of the Göttingen Graduate School for Neurosciences, Biophysics, and Molecular Biosciences (GGNB). J.S. was supported by the MINEDUC-UA project, code ANT 1855, Chile. Tohru Yamamoto (Kagawa University) generously provided access to the MDGA1 and MDGA2 mutant mouse lines, and Ann Marie Craig (University of British Columbia) provided breeding pairs for both mouse lines. The authors are grateful to Miso Mitkovski and the MPI-NAT Light Microscopy Facility for assistance with analysis of immunolabeling images, and to Fritz Benseler, the AGCT Lab, and the MPI-NAT Animal Facility for excellent technical support.

## Author contribution

D.K.-B. and N.B. conceived the current study. D.K-B. and H.A. designed immunohistochemical experiments, and H.A. and S.W. performed and analyzed immunohistochemical data with assistance from T.Z. and J.S.. H.T. and T.Z. designed brain slice electrophysiology experiments, and T.Z. conducted slice electrophysiological recordings with assistance from H.T., F.J.L.-M. E.G. performed pilot experiments with the help of J.S.R. H.T. provided custom software routines for data analysis. D.K.-B. designed behavioral experiments, and S.W. performed behavioral experiments with assistance from D.K.-B. and J.S.. N.B., J.W. and M.S. provided funding and equipment. D.K.-B., T.Z. and H.A. wrote the manuscript, N.B. and H.T. edited the manuscript, and all authors approved its final version.

