## Supplementary Information for "Functional Neuroligin-2-MDGA1 interactions differentially regulate synaptic GABA_A_Rs and cytosolic gephyrin aggregation"

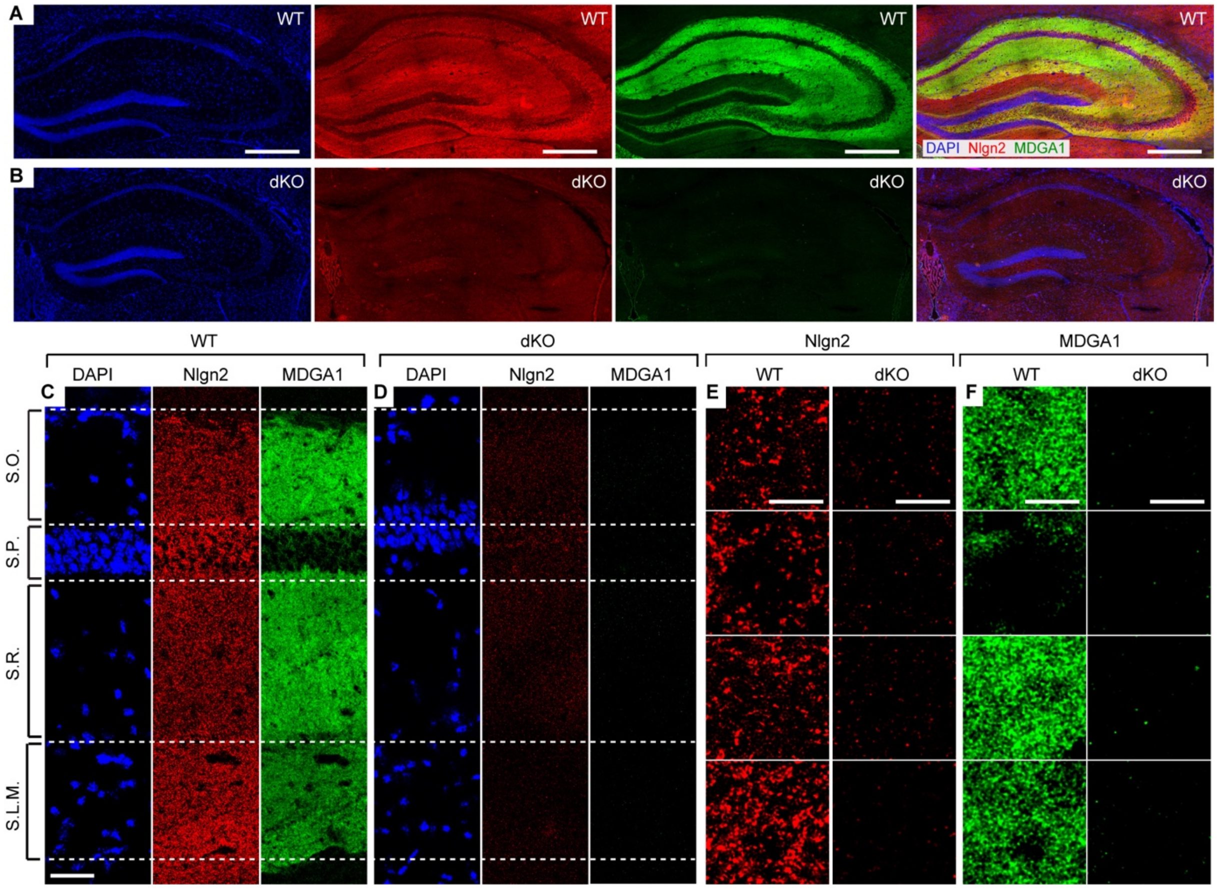


**Supplementary Figure 1. Validation of the specificity of antibodies against MDGA1 and Nlgn2.** (A-B) Photomicrographs showing an overview of the hippocampus in WT (A) versus Nlgn2-MDGA1 dKO mice (B) labelled with DAPI (blue), and antibodies against Nlgn2 (red) and MDGA1 (green). Scale bar 500 µm. (C-D) Photomicrographs showing an overview of area CA1 labelled with DAPI, and with antibodies against Nlgn2 and MDGA1 in WT (C) versus Nlgn2 / MDGA1 dKO mice (D). Scale bar 50 µm. (E-F) High magnification photomicrographs showing Nlgn2 and MDGA1 labeling within different hippocampal layers in WT (E) versus Nlgn2 / MDGA1 dKO mice (F), scale bar 5 µm.

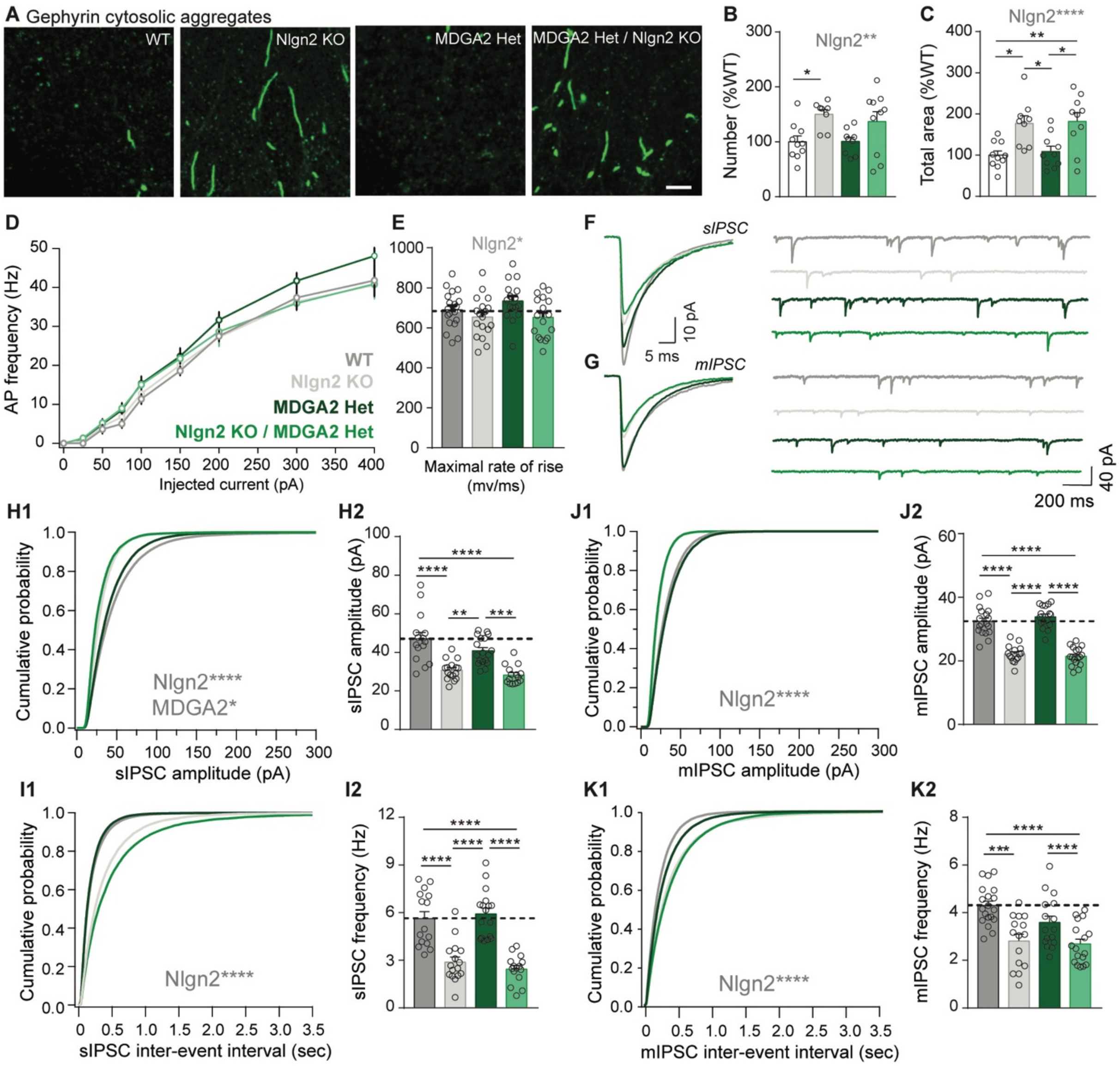


**Supplementary Figure 2. Heterozygous MDGA2 deletion shows no effects on the formation of gephyrin aggregates and on GABAergic transmission in CA1 pyramidal cells.** (A) High magnification photomicrographs of gephyrin aggregates in WT, Nlgn2 KO, MDGA2 Het and Nlgn2 KO / MDGA2 Het mice. Scale bar 5 µm. (B-C) Quantification of the number (B) and the total area (C) of gephyrin aggregates, expressed as percentage of WT. Statistically significant ANOVA comparisons are marked in gray at the top of panels and listed in Supplementary Table 3. For all other ANOVA comparisons, F<1. Post-hoc analysis (Tukey's comparison test): * p<0.05, ** p<0.01, *** p<0.001, **** p<0.0001. Error bars represent SEM, and each circle represents an experimental animal (n = 8-10). (D) Frequency of action potentials (APs) in response to steps of injected current. (E) Quantification of the maximal rate of AP rise in WT, Nlgn2 KO, MDGA2 Het, Nlgn2 KO / MDGA2 Het mice. (F) Representative average sIPSCs and sIPSC recording traces from all four genotypes analysed. (G) Representative average mIPSCs and mIPSC recording traces from all four genotypes analysed. (H-I) Average cumulative distribution and bar-graphs showing the quantification of sIPSC amplitude (H) and sIPSC inter-event frequency (I). (J-K) Average cumulative distribution and bar-graphs showing the quantification of mIPSC amplitude (J) and mIPSC inter-event frequency (K). Statistically significant ANOVA comparisons are marked in gray at the top of panels and listed in Supplemental Table 3. For all other ANOVA comparisons, F<1. Post-hoc analysis (Tukey's comparison test): * p<0.05, ** p<0.01, *** p<0.001, **** p<0.0001. In bar graphs, each circle represents a single cell (n = 14-19 cells for APs and rate of rise; 16-18 cells for sIPSC recordings; 15-19 cells for mIPSC recordings; each from four animals per genotype).

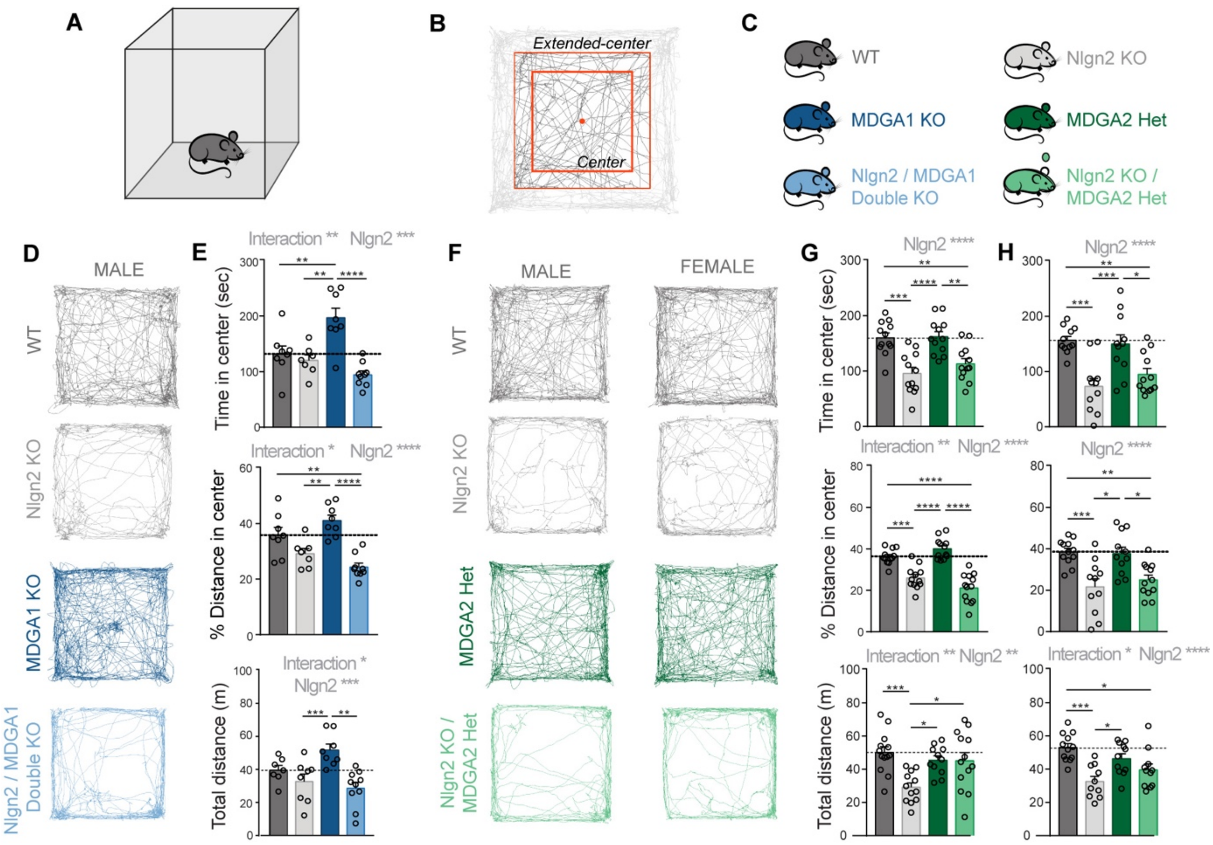


**Supplementary Figure 3. Heterozygous MDGA2 deletion does not alter anxiety-related avoidance behavior in Nlgn2 KO mice.** (A-C) Schematics representing the OF arena (A), the center (B), and the genotypes analyzed (C). (D) Representative tracks of OF exploration in MDGA1 male mice. (E) OF scores of MDGA1 male mice: Time spent in the anxiogenic region (top) of the OF arena, distance traveled in the center of the OF expressed as percentage of total distance traveled (center), total distance travelled in the OF (bottom). (F) Representative tracks of OF exploration in MDGA2 mice. (G) OF scores of MDGA2 male mice: Time spent in the anxiogenic region (top) of the OF arena, distance traveled in the center of the OF expressed as percentage of total distance traveled (center), total distance travelled in the OF (bottom). (H) OF scores of MDGA2 female mice: Time spent in the anxiogenic region (top) of the OF arena, distance traveled in the center of the OF expressed as percentage of total distance traveled (center), total distance travelled in the OF (bottom). Statistically significant ANOVA comparisons are marked in gray at the top of panels and listed in Supplementary Table 3. For all other ANOVA comparisons, F<1. Post-hoc analysis (Tukey's comparison test): * p<0.05, ** p<0.01, *** p<0.001, **** p<0.0001. Error bars represent SEM, and each circle represents an experimental animal (n = 7-9 for male MDGA1 set, n = 11-13 for female MDGA2 set, n = 10-12 for male MDGA2 set).

**Supplementary Table 1.** Analysis of the number and size of gephyrin, $GABA_AR\gamma2$ and VIAAT puncta in layers S.O. and S.L.M. of hippocampal area CA1 in WT, Nlgn2 KO, MDGA1 KO and Nlgn2 / MDGA1 dKO mice (all data expressed as %WT).

| | | WT | | Nlgn2 KO | | MDGA1 KO | | Nlgn2 / MDGA1 dKO | | Main source of variation | |
|---|---|---|---|---|---|---|---|---|---|---|---|
| | | n | Mean ± SEM | n | Mean ± SEM | n | Mean ± SEM | n | Mean ± SEM | F-value | p-value |
| Stratum Oriens (S.O.) | Gephyrin (number) | 8 | 100.00 ± 6.37 | 8 | 78.96 ± 6.96 | 8 | 86.56 ± 5.67 | 8 | 76.63 ± 9.63 | Nlgn2: $F_{(1,28)} = 4.484$ | 0.043 |
| | Gephyrin (size) | 8 | 100.00 ± 3.21 | 8 | 90.26 ± 3.28 | 7 | 94.27 ± 2.51 | 7 | 89.39 ± 3.27 | Nlgn2: $F_{(1,28)} = 4.227$ | 0.049 |
| | $GABA_AR\gamma2$ (number) | 8 | 100.00 ± 5.35 | 8 | 90.33 ± 6.30 | 9 | 93.14 ± 7.77 | 9 | 82.92 ± 6.54 | / | / |
| | $GABA_AR\gamma2$ (size) | 9 | 100.00 ± 3.91 | 8 | 81.46 ± 4.09 | 9 | 84.87 ± 3.48 | 9 | 81.99 ± 3.31 | Nlgn2: $F_{(1,31)} = 8.38$ Interaction $F_{(1,31)} = 4.48$ | Nlgn2: 0.007 Interaction 0.042 |
| | VIAAT (number) | 8 | 100.00 ± 15.71 | 8 | 101.19 ± 12.14 | 7 | 81.78 ± 12.35 | 7 | 74.33 ± 14.10 | / | / |
| | VIAAT (size) | 8 | 100.00 ± 2.44 | 8 | 98.99 ± 3.21 | 8 | 89.01 ± 3.09 | 8 | 87.57 ± 1.92 | MDGA1: $F_{(1,28)} = 18.4$ | <0.001 |
| Stratum lacunosum moleculare (SLM) | Gephyrin (number) | 7 | 100.00 ± 7.90 | 7 | 98.50 ± 8.66 | 7 | 109.89 ± 4.36 | 7 | 93.18 ± 2.79 | / | / |
| | Gephyrin (size) | 8 | 100.00 ± 3.83 | 8 | 95.64 ± 2.30 | 7 | 97.97 ± 1.76 | 7 | 88.85 ± 3.58 | / | / |
| | $GABA_AR\gamma2$ (number) | 8 | 100.00 ± 6.04 | 8 | 94.59 ± 4.05 | 8 | 107.69 ± 3.97 | 9 | 102.60 ± 4.38 | | |
| | $GABA_AR\gamma2$ (size) | 8 | 100.00 ± 4.27 | 9 | 85.47 ± 4.1 | 8 | 84.19 ± 2.74 | 9 | 83.23 ± 3.39 | Nlgn2: $F_{(1,30)} = 4.38$ MDGA1: $F_{(1,30)} = 5.94$ | Nlgn2: 0.045 MDGA1: 0.021 |
| | VIAAT (number) | 6 | 100.00 ± 26.44 | 6 | 106.59 ± 22.35 | 4 | 117.51 ± 23.79 | 4 | 120.34 ± 17.85 | / | / |
| | VIAAT (size) | 7 | 100.00 ± 3.32 | 6 | 100.82 ± 5.50 | 4 | 84.13 ± 3.68 | 4 | 95.16 ± 3.82 | MDGA1: $F_{(1,19)} = 7.09$ | MDGA1: 0.015 |

**Supplementary Table 2 (Part 1).** Analysis of the number and size of gephyrin, $GABA_AR\gamma2$ and VIAAT puncta in layers S.O., S.P., S.R. and S.L.M. of hippocampal area CA1 in WT, Nlgn2 KO, MDGA2 Het and Nlgn2 KO / MDGA2 Het mice (all data expressed as %WT).

| | | WT | | Nlgn2 KO | | MDGA2 Het | | Nlgn2 KO / MDGA2 Het | | Main source of variation | |
|---|---|---|---|---|---|---|---|---|---|---|---|
| | | n | Mean ± SEM | n | Mean ± SEM | n | Mean ± SEM | n | Mean ± SEM | F-value | p-value |
| Stratum Oriens (S.O.) | Gephyrin (number) | 11 | 100.00 ± 8.92 | 10 | 86.87 ± 6.49 | 11 | 104.31 ± 8.61 | 9 | 78.32 ± 6.99 | Nlgn2: $F_{(1,37)} = 5.951$ | Nlgn2: 0.020 |
| | Gephyrin (size) | 10 | 100.00 ± 3.03 | 10 | 91.35 ± 2.43 | 10 | 96.68 ± 2.77 | 10 | 94.13 ± 2.28 | Nlgn2: $F_{(1,37)} = 4.527$ | Nlgn2: 0,040 |
| | $GABA_AR\gamma2$ (number) | 12 | 100.00 ± 7.65 | 11 | 87.60 ± 8.94 | 12 | 117.91 ± 13.8 | 12 | 60.32 ± 8.05 | Interaction: $F_{(1,43)} = 5.130$<br>Nlgn2: $F_{(1,43)} = 12.31$ | Interaction: 0.029<br>Nlgn2: 0.001 |
| | $GABA_AR\gamma2$ (size) | 10 | 100.00 ± 2.55 | 10 | 86.17 ± 3.14 | 12 | 93.84 ± 5.02 | 11 | 76.82 ± 2.17 | Nlgn2: $F_{(1,39)} = 18.65$<br>MDGA2: $F_{(1,39)} = 4.71$ | Nlgn2: <0.001<br>MDGA2: 0.036 |
| | VIAAT (number) | 7 | 100.00 ± 17.70 | 7 | 84.99 ± 10.96 | 6 | 8297 ± 18.43 | 6 | 58.58 ± 9.60 | / | / |
| | VIAAT (size) | 7 | 100.00 ± 5.32 | 7 | 99.28 ± 3.09 | 6 | 89.99 ± 2.47 | 7 | 93.22 ± 6.13 | / | / |
| Stratum Pyramidale (S.P.) | Gephyrin (number) | 9 | 100.00 ± 10.72 | 9 | 88.69 ± 6.88 | 9 | 79.64 ± 7.10 | 9 | 70.07 ± 8.67 | MDGA2: $F_{(1,32)} = 5.281$ | MDGA2: 0.028 |
| | Gephyrin (size) | 9 | 100.00 ± 2.40 | 9 | 90.28 ± 3.64 | 9 | 92.68 ± 3.08 | 8 | 89.18 ± 1.27 | Nlgn2: $F_{(1,31)} = 5.515$ | Nlgn2: 0.025 |
| | $GABA_AR\gamma2$ (number) | 9 | 100.00 ± 3.98 | 10 | 81.33 ± 7.56 | 10 | 84.78 ± 8.66 | 9 | 59.50 ± 5.25 | Nlgn2: $F_{(1,34)} = 10.36$<br>MDGA2: $F_{(1,34)} = 7.359$ | Nlgn2: 0.003<br>MDGA2: 0.010 |
| | $GABA_AR\gamma2$ (size) | 10 | 100.00 ± 5.39 | 10 | 88.88 ± 4.56 | 10 | 87.27 ± 3.88 | 10 | 70.77 ± 3.43 | Nlgn2: $F_{(1,36)} = 10.00$<br>MDGA2: $F_{(1,36)} = 12.34$ | Nlgn2: 0.003<br>MDGA2: 0.001 |
| | VIAAT (number) | 8 | 100.00 ± 19.34 | 8 | 91.39 ± 10.93 | 8 | 71.34 ± 11.78 | 7 | 86.27 ± 12.51 | / | / |
| | VIAAT (size) | 8 | 100.00 ± 9.21 | 8 | 98.07 ± 2.99 | 7 | 109.34 ± 3.65 | 7 | 103.67 ± 4.69 | / | / |
| Stratum Radiatum (S.R.) | Gephyrin (number) | 11 | 100.00 ± 10.69 | 9 | 85.62 ± 8.19 | 11 | 94.89 ± 9.10 | 8 | 92.60 ± 3.56 | / | / |
| | Gephyrin (size) | 10 | 100.00 ± 3.27 | 9 | 95.68 ± 1.80 | 9 | 94.87 ± 2.51 | 10 | 93.62 ± 2.30 | / | / |
| | $GABA_AR\gamma2$ (number) | 11 | 100.00 ± 8.02 | 11 | 75.02 ± 9.24 | 12 | 90.91 ± 9.06 | 12 | 55.97 ± 8.86 | Nlgn2: $F_{(1\ 42)} = 11.48$ | Nlgn2: 0.002 |
| | $GABA_AR\gamma2$ (size) | 10 | 100.00 ± 3.37 | 10 | 83.94 ± 2.98 | 11 | 99.99 ± 5.22 | 11 | 78.23 ± 3.04 | Nlgn2: $F_{(1,38)} = 24.41$ | Nlgn2: <0.001 |
| | VIAAT (number) | 7 | 100.00 ± 25.15 | 7 | 81.17 ± 13.19 | 7 | 68.68 ± 11.52 | 5 | 90.43 ± 7.13 | / | / |
| | VIAAT (size) | 7 | 100.00 ±7.12 | 7 | 101.76 ± 2.85 | 7 | 92.45 ± 4.57 | 7 | 99.49 ± 4.95 | / | / |

**Supplementary Table 2 (Part 2).** Analysis of the number and size of gephyrin, GABA$_A$Rγ2 and VIAAT puncta in layers S.O., S.P., S.R. and S.L.M. of hippocampal area CA1 in WT, Nlgn2 KO, MDGA2 Het and Nlgn2 KO / MDGA2 Het mice (all data expressed as %WT).

| | | WT | | Nlgn2 KO | | MDGA2 Het | | Nlgn2 KO / MDGA2 Het | | Main source of variation | |
|---|---|---|---|---|---|---|---|---|---|---|---|
| | | n | Mean ± SEM | n | Mean ± SEM | n | Mean ± SEM | n | Mean ± SEM | F-value | p-value |
| Stratum lacunosum moleculare (S.L.M.) | Gephyrin (number) | 10 | 100 .00 ±11.18 | 9 | 99.21 ± 3.61 | 11 | 88.76 ± 5.43 | 8 | 108.41 ± 6.41 | / | / |
| | Gephyrin (size) | 11 | 100.00 ± 3.60 | 9 | 94.38 ± 1.26 | 11 | 93.48 ± 2.56 | 10 | 92.63 ± 1.90 | / | / |
| | GABA$_A$Rγ2 (number) | 9 | 100.00 ± 8.93 | 7 | 85.92 ± 3.43 | 10 | 82.21 ± 9.79 | 9 | 71.60 ± 9.97 | / | / |
| | GABA$_A$Rγ2 (size) | 10 | 100.00 ± 4.8 | 9 | 84.29 ± 2.72 | 9 | 88.32 ± 3.23 | 9 | 77.98 ±1.69 | Nlgn2: $F_{(1,33)} = 14.50$<br>MDGA2: $F_{(1,33)} = 6.914$ | Nlgn2: <0.001<br>MDGA2: 0.013 |
| | VIAAT (number) | 5 | 100.00 ± 20.59 | 6 | 49.13 ± 11.20 | 8 | 30.52 ± 8.52 | 7 | 32.86 ± 6.18 | Interaction: $F_{(1,22)} = 5.483$<br>Nlgn2: $F_{(1,22)} = 4.549$<br>MDGA2: $F_{(1,22)} = 14.22$ | Interaction: 0.029<br>Nlgn2: 0.044<br>MDGA2: 0.001 |
| | VIAAT (size) | 7 | 100.00 ± 7.76 | 6 | 101.08 ± 10.61 | 8 | 87.57 ± 7.34 | 7 | 99.64 ± 3.74 | / | / |

**Supplementary Table 3.** Two-way ANOVA comparisons for Supplementary Figures 2-3.

| Figure | Nlgn2 x MDGA2 interaction | | Main effect of Nlgn2 | | Main effect of MDGA2 | |
|---|---|---|---|---|---|---|
| | F-value | p-value | F-value | p-value | F-value | p-value |
| **S2 B** | $F_{(1,35)} < 1$ | 0.913 | $F_{(1,35)} = 21.46$ | <0.0001 | $F_{(1, 5)} < 1$ | 0.662 |
| **S2 C** | $F_{(1,33)} < 1$ | 0.582 | $F_{(1,33)} = 11.73$ | 0.0017 | $F_{(1,33)} < 1$ | 0.621 |
| **S2 F** | $F_{(1,66)} < 1$ | 0.338 | $F_{(1,66)} = 6.247$ | 0.0149 | $F_{(1,66)} < 1$ | 0.382 |
| **S2 I** | $F_{(1,59)} = 1.095$ | 0.300 | $F_{(1,59)} = 78.98$ | <0.0001 | $F_{(1,59)} < 1$ | 0.843 |
| **S2 J** | $F_{(1,60)} < 1$ | 0.359 | $F_{(1,60)} = 51.10$ | <0.0001 | $F_{(1,60)} = 4.661$ | 0.035 |
| **S2 K** | $F_{(1,62)} = 1.655$ | 0.203 | $F_{(1,62)} = 171.6$ | <0.0001 | $F_{(1,62)} < 1$ | 0.679 |
| **S2 L** | $F_{(1,63)} = 1.590$ | 0.212 | $F_{(1,63)} = 26.12$ | <0.0001 | $F_{(1,63)} = 3.204$ | 0.078 |
| **S3G** ***Time in centre*** | $F_{(1,44)} < 1$ | 0.777 | $F_{(1,44)} = 11.61$ | 0.0014 | $F_{(1,44)} = 1.727$ | 0.196 |
| **S3G** ***%Distance in centre*** | $F_{(1,44)} = 5.679$ | 0.022 | $F_{(1,44)} = 56.75$ | <0.0001 | $F_{(1,44)} < 1$ | 0.401 |
| **S3G** ***Total distance*** | $F_{(1\ 45)} = 7.953$ | 0.007 | $F_{(1,45)} = 8.532$ | 0.0054 | $F_{(1,45)} = 2.383$ | 0.130 |
| **S3H** ***Time in centre*** | $F_{(1,41)} = 1.350$ | 0.252 | $F_{(1,41)} = 13.21$ | 0.0008 | $F_{(1,41)} < 1$ | 0.707 |
| **S3H** ***%Distance in centre*** | $F_{(1,41)} < 1$ | 0.961 | F($_{1, 41)}$ = 20.41 | <0.0001 | $F_{(1,41)} < 1$ | 0.822 |
| **S3H** ***Total distance*** | $F_{(1,40)} = 5.186$ | 0.028 | $F_{(1,40)} = 20.65$ | <0.0001 | $F_{(1,40)} < 1$ | 0.959 |
| **Figure** | **Nlgn2 x MDGA1 interaction** | | **Main effect of Nlgn2** | | **Main effect of MDGA1** | |
| | **F-value** | **p-value** | **F-value** | **p-value** | **F-value** | **p-value** |
| **S3E** ***Time in centre*** | $F_{(1,27)} = 2.473$ | 0.128 | $F_{(1,27)} = 21.55$ | <0.001 | $F_{(1,27)} = 3.264$ | 0.082 |
| **S3E** ***%Distance in centre*** | $F_{(1,29)} = 2.814$ | 0.104 | $F_{(1,29)} = 28.64$ | <0.001 | $F_{(1,29)} < 1$ | 0.832 |
| **S3E** ***Total distance*** | $F_{(1,29)} = 4.665$ | 0.039 | $F_{(1,29)} = 16.04$ | <0.001 | $F_{(1,29)} = 1.198$ | 0.283 |

**Supplementary Table 4.** Passive and AP properties of CA1 pyramidal cells in WT, Nlgn2 KO, MDGA2 Het and Nlgn2 KO / MDGA2 Het mice.

| | **WT** | | **Nlgn2 KO** | | **MDGA2 Het** | | **Nlgn2 KO / MDGA2 Het** | | **Main source of variation** | |
|---|---|---|---|---|---|---|---|---|---|---|
| | n | Mean ± SEM | n | Mean ± SEM | n | Mean ± SEM | n | Mean ± SEM | F-value | p-value |
| **Membrane resistance (MOhm)** | 41 | 92.07 ± 4.41 | 35 | 97.43 ± 3.06 | 36 | 99.59 ± 5.26 | 35 | 104.19 ± 4.64 | \ | \ |
| **Membrane capacitance, proximal compartments (pF)** | 41 | 45.67 ± 1.94 | 35 | 45.48 ± 2.03 | 36 | 40.53 ± 1.39 | 35 | 42.65 ± 2.34 | MDGA2: $F_{(1,143)} = 4.14$ | 0.044 |
| **Membrane capacitance, distal compartments (pF)** | 41 | 116.42 ± 4.95 | 35 | 106.48 ± 3.75 | 36 | 107.6 ± 5.55 | 35 | 109.56 ± 4.85 | \ | \ |
| **Resting membrane potential (mV)** | 20 | -58.83 ± 1.19 | 19 | -60.39 ± 1.46 | 17 | -56.99 ± 1.75 | 17 | -55.92 ± 1.85 | MDGA2: $F_{(1,69)} = 4.47$ | 0.038 |
| **AP threshold (mV)** | 19 | -45.56 ± 1.03 | 18 | -45.50 ± 0.77 | 17 | -45.58 ± 0.77 | 17 | -45.65 ± 0.55 | \ | \ |
| **AP amplitude (mV)** | 19 | 117.16 ± 0.98 | 18 | 119.14 ± 1.47 | 17 | 120.34 ± 1.11 | 17 | 116.89 ± 1.46 | Interaction: $F_{(1,67)} = 4.60$ | 0.036 |
| **AP Maximal rate of rise (mv/ms)** | 19 | 690.66 ± 21.03 | 18 | 654.42 ± 24.01 | 16 | 734.23 ± 24.52 | 17 | 652.39 ± 24.99 | Nlgn2: $F_{(1,66)} = 6.25$ | 0.015 |
